# A diffusion-like process enables expansion of advantaged gene mutations in human colonic epithelium

**DOI:** 10.1101/2020.07.10.193748

**Authors:** Cora Olpe, Doran Khamis, Maria Chukanova, Richard Kemp, Kate Marks, Cerys Tatton, Cecilia Lindskog, Nefeli Skoufou-Papoutsaki, Anna Nicholson, Roxanne Brunton-Sim, Shalini Malhotra, Rogier ten Hoopen, Rachel Stanley, Doug Winton, Edward Morrissey

## Abstract

Colorectal cancer is thought to arise when the mutational burden of the clonal population of stem cells within a colonic crypt exceeds a certain threshold. Therefore, quantification of the fixation and subsequent expansion of somatic mutations in histologically normal epithelium is key to understanding colorectal cancer initiation. Here, using immunohistochemistry, loss of the histone demethylase KDM6A in normal human colonic epithelium is visualised. Interpretation of the age-related behaviour of KDM6A-negative clones revealed significant competitive advantage in intra-crypt dynamics. Further, subsequent clonal expansion into multi-crypt patches was quantified to reveal a significant 5-fold increase in crypt fission rate. To accomodate the local accumulation of new crypts, the role of crypt fusion was considered. However, no compensatory increase in fusion rate was found. Instead, evidence for crypt diffusion is presented and proposed as a means of accommodating clonal expansions. The threshold fission rate at which diffusion fails to accommodate new crypts, and which may promote polyp growth, is defined.

## Introduction

The development of epithelial cancers is defined by mutations that become fixed and expand within host tissues. In human colon somatic variants become fixed only if they outcompete wildtype neighbours to successfully populate entire colonic crypts (1). The subsequent process of expansion is mediated by a crypt replication process, termed fission (2–4). Where clones can be visualised in situ, fission rates can be directly calculated from changes in clone size distributions with age (5–11). These studies confirm a low homeostatic fission rate in adult colon. In contrast, the development of focal neoplastic disease is thought to be a consequence of elevated fission rates. Most notably loss of the tumour suppressor gene APC is thought to generate adenomas in this way (12–15). However, recently we demonstrated that crypts with increased fission rates, due to loss of the X-linked gene STAG2, could generate large expansions that appear phenotypically normal (1).

More widely it is recognised that many of the renewing epithelia, including colon, acquire a substantial burden of cancer driver mutations while also remaining apparently normal (16–18). Clonal expansions without an overt phenotype but predisposing to oncogenic transformation may explain the recent interpretation that colorectal cancers arise as a single expansion event when a combination of factors reaches a critical threshold (19). This suggests that understanding the development of cancer requires an appreciation not only of how cell and tissue level processes allow mutational burden to be achieved, but also how it is configured within small or large expansions.

There does not appear to be an increase in the net density of crypts, or of colonic epithelial area, with age (20). This raises the question: how are local clonal expansions arising from elevated fission rates accommodated? One explanation might lie in crypt fusion. This process has recently been described in the mouse small intestine and human colon and could counteract the consequences of fission (21, 22). However, it remains unclear if fusion is a stochastic process occurring independently of fission or if they are locally co-regulated. The latter possibility may be particularly relevant to pro-oncogenic mutations as fusions at the edge of mutant patches could allow an effective local invasion of wildtype crypts with a high probability that wildtype cells will subsequently be displaced.

KDM6A (UTX) is an X-linked gene encoding a histone demethylase that specifically targets di- and tri-methyl groups on lysine 27 of histone H3 (H3K27me2/3). Inactivating mutations and deletions of KDM6A have been identified in a variety of human cancers including multiple myeloma, colon, bladder, prostate, medulloblastoma, oesophageal and renal cancer (23–26). KDM6A is one of the 127 significantly mutated genes in The Cancer Genome Atlas (TCGA) study that analysed 3281 tumours derived from 12 cancer types (27).

Here, in seeking additional cancer driver events that can be spatially visualised as somatic clones, we identify mutations of KDM6A as possessing advantage in both intra-crypt fixation and subsequent expansion. The large multicrypt clones resulting from elevated rates of crypt fission are investigated to study the impact of expansion on crypt packing and the role of crypt fusion in relieving overcrowding. The increased fission rate within KDM6A^−^ clones is not accompanied by an increase in crypt fusion, suggesting the two processes are driven by independent mechanisms and that fusion does not act to relieve local overcrowding. Instead, it is proposed that new crypts generated by fission can be accommodated by localised crypt diffusion up to a threshold beyond which hyperplastic and neoplastic lesions may form.

## Results

### KDM6A-negative clones are advantaged in stem cell competition

We have previously shown that visualisation of loss of X-linked genes can be used as clonal marks to quantify human colonic stem cell dynamics (1). In attempting to expand this methodology to X-linked genes with cancer association, clonal loss of KDM6A was identified by immunohistochemistry with two independent antibodies on normal human colonic epithelia (Figure 1A & B).

**Figure 1.**
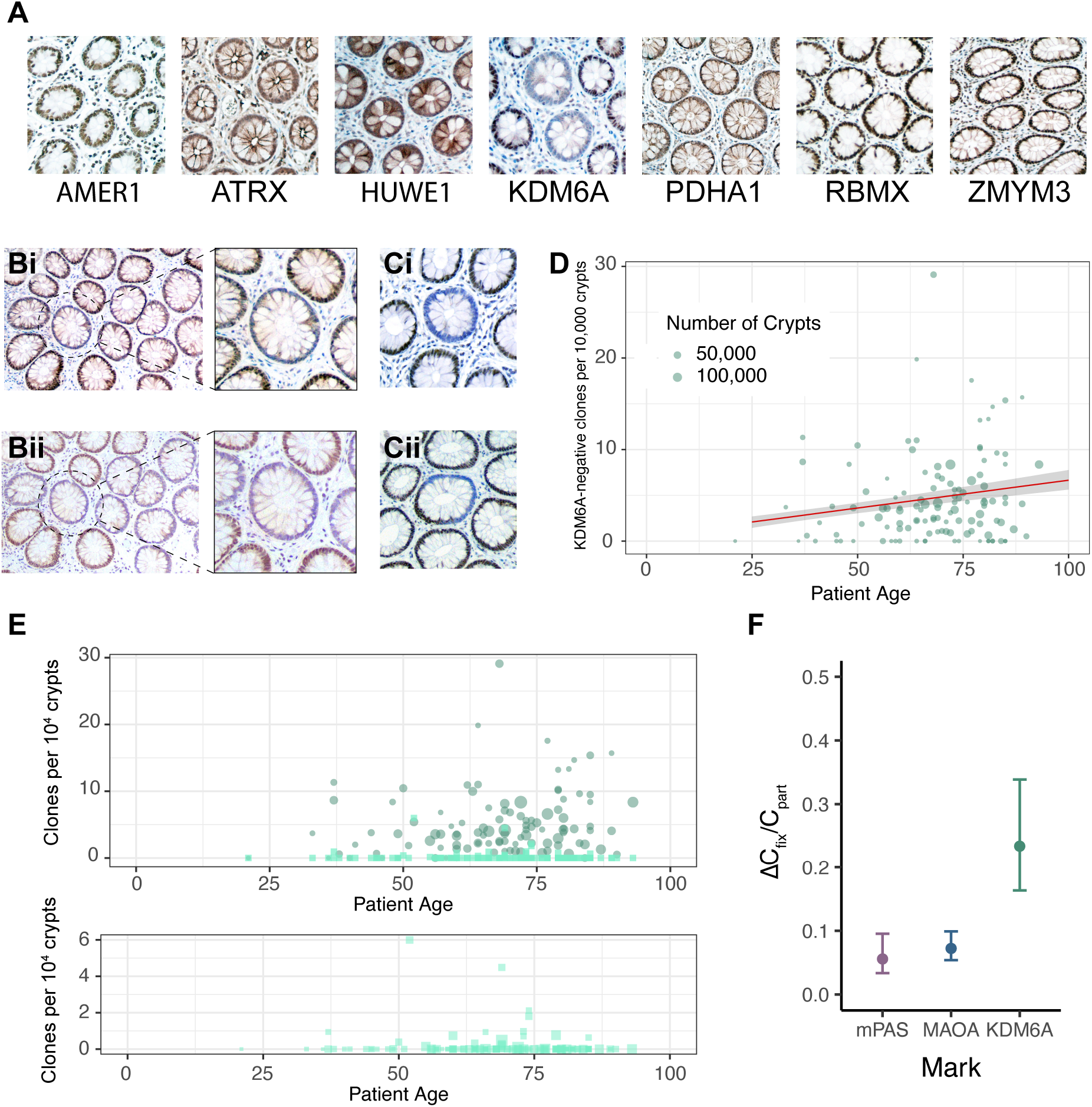
Loss of KDM6A results in advantaged stem cell dynamics. (A) Representative images of IHC for cancer-associated X-linked genes on human colonic sections. Out of the 7 assessed genes, a staining pattern indicative of loss-of-function mutation was only found for KDM6A (middle panel). (B) Serial human colonic sections stained with two independent antibodies against KDM6A: (i) human protein atlas, (ii) Cell Signaling Technology. KDM6A^−^ crypts are circled and enlarged. (C) Representative images of KDM6A^−^ WPC (i) and PPC (ii). (D) Regression analysis (red line) showing accumulation of WPC (ΔC_*fix*_: 6.04 × 10^−6^ per year) and 95% confidence interval in grey. (E) Top panel: Frequency plot showing age-related behaviours of KDM6A-negative WPC (circles, darker) and PPC (squares, lighter). Bottom panel showing PPC only on expanded y-axis. (F) Plot showing increased ratio of ΔC_fix_/C_part_ for KDM6A (0.23, 95% CI: 0.16 –0.34) as compared to the neutral markers mPAS and MAOA (replotted from (1)). Error bars = 95% CI.

Intra-crypt dynamics that describe the accumulation of clones wholly populating entire crypts (WPC) from partly populated (PPC) transition forms were determined for KDM6A^−^ clones as previously described (1). Colonic FFPE sections from 120 patients aged 21–93 years were stained to visualise KDM6A^−^ clones. The total number of crypts per *en face* section was determined using a neural-network based image analysis tool (DeCryptICS, https://github.com/MorrisseyLab/DeCryptICS) and KDM6A^−^ clones manually counted (Figure 1C). Regression analysis revealed an age-related increase in the frequency of WPC with slope, ΔC_*fix*_, of 6.0 × 10^−6^ KDM6A^−^ clones per year (95% CI: 4.7-7.8 × 10^−6^) (Figure 1D). As expected, the frequency of PPC, C_*part*_, remained constant at all ages [2.6 × 10^−5^ (95%CI: 1.9 – 3.4 × 10^−5^)] (Figure 1E). The ratio between these two values provides a means of comparing the effect of different gene-specific mutations on stem cell dynamics that is independent of the mutation rate (1). The value obtained for ΔC_*fix*_/C_part_ was 0.23 (95% CI: 0.16– 0.34) around 4.5-fold higher than that for neutral marks and indicates a competitive advantage (Figure 1F). The advantage conferred by KDM6A loss was expressed as a probability of replacement (P_R_) at each round of stem cell replacement whereby values above 0.5 (neutral probability of loss or gain of a mutant stem cell) identify positive bias. The P_R_ for KDM6A was 0.76 (95% CI: 0.60–1.0).

### KDM6A-negative clones expand by 5-fold increased crypt fission

Expansion of individual KDM6A^−^ clones was recognisable as large patches that were frequently comprised of more than 10 crypts (Figure 2A). To confirm the clonal origin of such patches, we used laser capture dissection followed by targeted sequencing that covered 3.6 kb of exonic and flanking intronic sequence of *KDM6A* (24 amplicons), including sites frequently mutated in human cancers (Figure 2B & Sup. Figure 1). Amplicon libraries from four patches failed QC. Of the remaining seven patches mutations were identified in four at mutant allele frequencies consistent with patient sex and estimated stromal fraction within the captured material (Figure 2B and Supplemental Table 1). Of note, the mutation found in intron 18 was present in two independent patches. Furthermore, the mutations S1154* and W1193* have previously been found in cancer samples (COSMIC database). These results both confirm antibody specificity for KDM6A and the clonality of multicrypt patches that share only a single mutation.

**Figure 2.**
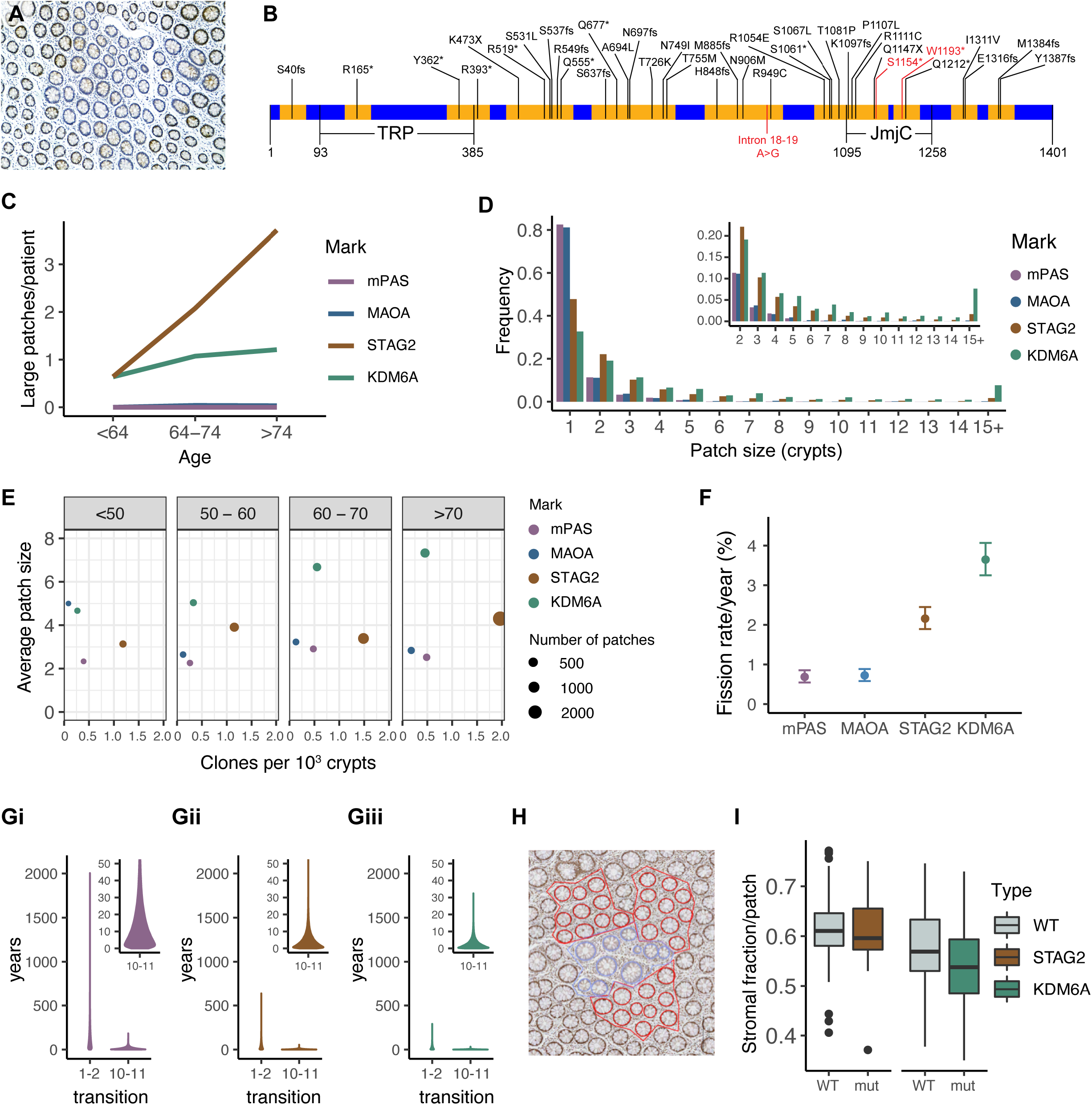
Molecular and phenotypic characterisation of KDM6A^−^ patches. (A) Representative image of large KDM6A-negative multicrypt patch. (B) Schematic shows KDM6A cDNA structure with areas covered by amplicons (yellow) and annotated with mutations found in COSMIC. Mutations identified in KDM6A-negative patches are indicated (red). (C) Plot showing mean number of large (≥10 crypts) patches per patient for age groups shown. (D) Histogram showing the frequency of different patch sizes for mPAS, MAOA, KDM6A and STAG2 across all ages. Inset shows patch size ≥2 crypts on expanded y-axis. (E) Dot plot of mean clone frequency plotted against mean average patch size for multicrypt patches in age groups shown for mPAS, MAOA, STAG2 and KDM6A. (F) Plot showing inferred fission rates/crypt/year for KDM6A compared to mPAS, MAOA and STAG2 (data from (1) replotted). Error bars = 95% CI. (G) Simulation data showing the time in years taken for transitions between patch sizes 1-2 and 10-11 for (i) mPAS, (ii) STAG2 and (iii) KDM6A. Insets show 10-11 transition on expanded y-axis. (H) Image showing patch selection for assessment of crypt packing density within KDM6A-(red) and adjacent wildtype crypts (blue). (I) Box plots showing fraction of patch stromal space for STAG2^−^ and KDM6A^−^negative mutant and adjacent wildtype patches comprising 10 crypts.

The age-related size distribution of multicrypt clones was analysed to infer the fission rate associated with KDM6A loss and to compare that to those previously described for neutral (MAOA and mPAS) and positively biased clonal marks (STAG2) (1). This revealed that the frequency of large clones (≥10 crypts/patch) increases with age for STAG2^−^ and KDM6A^−^ but not for MAOA and mPAS (Figure 2C). Loss of KDM6A generates a higher proportion of large patches than the other clonal marks while STAG2 loss results in more clones due to a higher event rate (Figure 2D & E). From these patch size distributions, the crypt fission rate associated with loss of KDM6A was calculated to be to 3.6% per year (95% CI: 3.2-4.1), approximately 5-fold higher than the background homeostatic rate previously derived from neutral clonal marks (Figure 2F). Consequently, in individuals over 80 years of age 13.5% of KDM6A^−^ clones are found as patches comprising more than 5 crypts compared to 4.8% for STAG2, 1.7% for mPAS and 0.8% for MAOA.

### KDM6A^−^ and STAG2^−^ patches lack significant local overcrowding

The probability of fission remains low even when elevated as for STAG2 and KDM6A. Many single crypt clones never expand. Clones forming from those that do are more likely to undergo subsequent fission as the probability scales with the number of crypts in each clone. Consequently, the time taken to increase the number of crypts in a specific area decreases rapidly with increasing patch size (Figure 2G). For example, mathematical modelling based on the inferred fission rate for KDM6A^−^ crypts indicates the median time taken to grow from 1 to 2 crypts is 19 years, but only 2 years to grow from 10 to 11. Therefore, recently formed larger patches might be expected to demonstrate overcrowding.

To test if larger patches are more densely packed, the area occupied by crypts and their surrounding stroma was determined for 24 STAG2^−^ and 20 KDM6A^−^ clones containing ten crypts (Figure 2H). The fraction of each patch occupied by stroma was then calculated. Adjacent to each mutant clone three random groups of 10 crypts were defined as control ‘patches’ and similarly analysed. Comparing the fraction of each patch occupied by stroma to adjacent wildtype groupings indicated a slight trend towards increased packing density for STAG2^−^ as well as KDM6A^−^ crypts that failed to reach significance (Figure 2I). Considering that a lack of overcrowding may stem from a decrease in crypt size, the areas of crypts were measured. This revealed that KDM6A^−^ crypt sections are around x1.3 the size of adjacent wildtype crypts (p-value <0.001). No difference was found between STAG2^−^ crypts and their wildtype neighbours (Supplemental Figure 2). Lack of overcrowding cannot be attributed to reduced crypt size for either STAG2 or KDM6A loss.

Together these observations suggest that crypts even within relatively recent clonal expansions avoid overcrowding to largely achieve ambient density.

### Evidence for crypt fusion

The lack of overcrowding in KDM6A^−^ clones suggests a mechanism counteracting the localised increase in fission. An opposing process of crypt fusion has been recognised in the mouse (21). A homeostatic human fusion rate has recently been estimated and proposed as a mechanism to relieve local strain (22). This estimate was based on a calculation of fission and deriving a fusion rate assuming equivalence in the rate of both processes. Clearly on a tissue wide basis such a balance of rates could act to maintain constant crypt density. However, local increases in fission rates in advantaged clonal expansions can only balance if fission and fusion are coordinately regulated by local cross talk.

We first sought confirmation that fusion occurs. The evidence in human epithelium is based on the presence of branched crypts within which one branch is marked by clonal loss of mitochondrial CCO activity while the other is not. These are interpreted as transition intermediates in an active crypt fusion process (22). Analysis of *en face* tissue sections stained to visualise clonal mPAS positivity as well as loss of STAG2 and KDM6A confirmed the existence of rare heterotypic branched forms in normal human colonic epithelium. Analysis of over 2×10^6^ crypts in sections from 80 individuals containing mPAS^+^ clones identified 32 candidate mPAS^+^ branched forms that were either mixed (mutant and wild type, M/W) or fully mutant (M/M) (Figure 3A). Of the 13 with mixed staining (M/W) the positive epithelium was always segregated into one branch and not the other.

**Figure 3.**
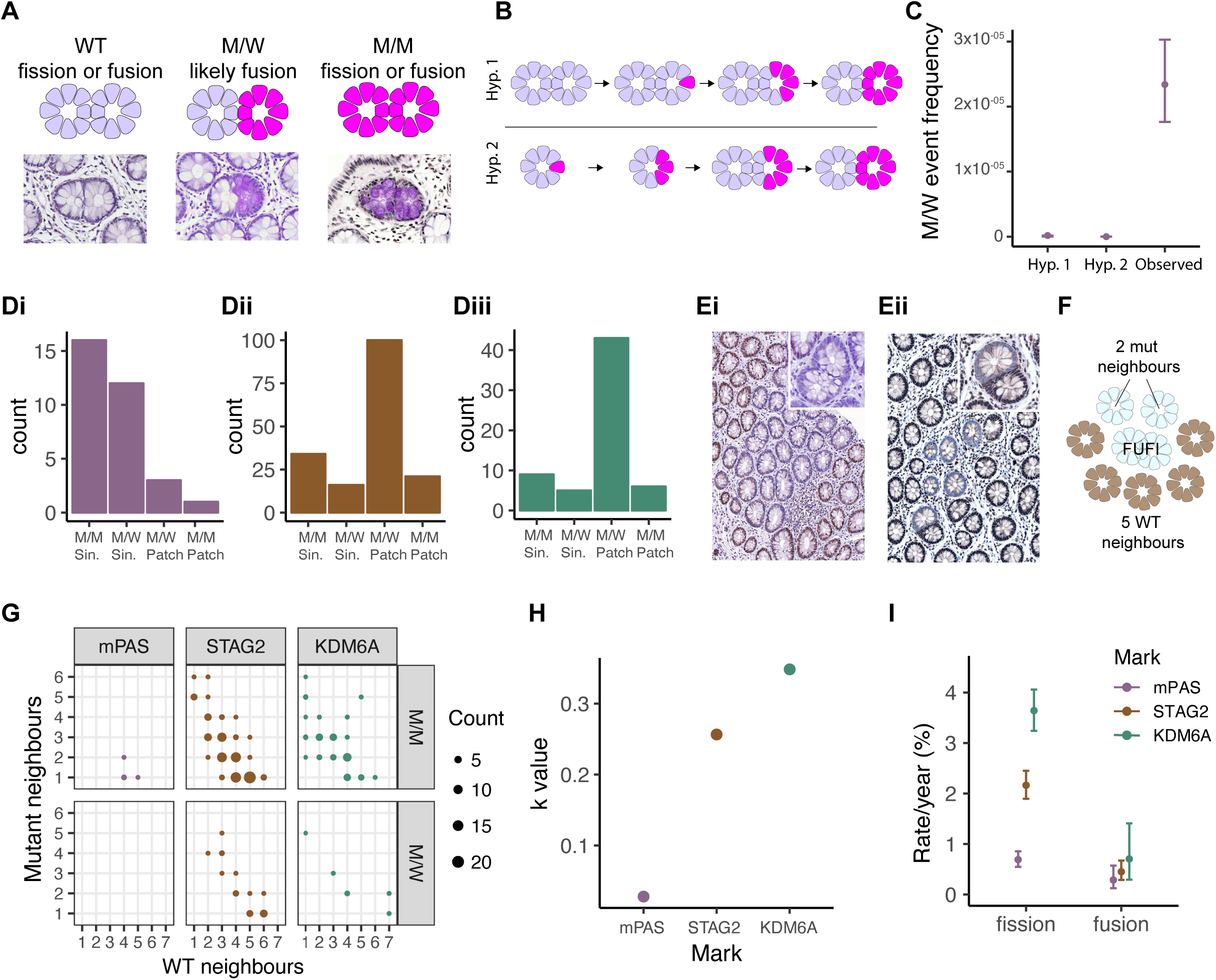
Crypt fission and fusion are independently regulated processes. (A) Schematic and representative mPAS-stained images of three types of fusion or fission forms. (B) Schematic representation of alternative origins of M/W forms. Top: Hypothesis 1 (Hyp1): stem cell mutation in one branch of a crypt undergoing fission followed by monoclonal coversion of that branch. Bottom: Hypothesis 2 (Hyp 2): fission of a pre-existing partially populated crypt with segregation of mutant and wild type epithelium into each branch, followed by monoclonal conversion. (C) Comparison of M/W event frequencies simulated for hypotheses described in (B) and the observed frequency. Error bars = 95% CI. (D) Bar graphs showing numbers of FUFI types scored for (i) mPAS, (ii) KDM6A, (iii) STAG2. (E) Representative images of FUFIs in patches: (i) KDM6A M/M FUFI at border of large patch; (ii) STAG2 M/W FUFI at patch border. Inset shows enlarged FUFI. (F) Schematic showing scoring of FUFI neighbours used for calculation of k. (G) Dot plot showing neighbouring crypt status of patch border FUFIs (either M/M or M/W type) for mPAS, STAG2 and KDM6A. Each dot represents one or more FuFis for which neighbours were scored. (H) Plot comparing the value of *k*, representing the fraction of M/M FUFIs that are fusions, for mPAS, STAG2 and KDM6A. (I) Plot comparing the derived fission and fusion rates associated with mPAS, STAG2 and KDM6A. mPAS and STAG2 fission rates correspond to data from (1), replotted. Error bars = 95% CI.

An alternative interpretation of branched crypts is that they are intermediate fission forms and that heterotypic staining arises due to mutations occurring or segregating into a single branch (Figure 3B). We formally considered this possibility using the fusion duration estimate derived by Baker and colleagues as well as our previous estimates of *de novo* mutation probability and clone fixation rates that together determine the frequency of monoclonal crypts present in individuals of different age (22, 28, 29). Within the relatively small number of branched crypts present none are predicted to contain monoclonal crypt branches by either mechanism (Figure 3C). This suggests that heterotypic branched crypts represent genuine intermediates participating in an active fusion process. Noting that the bulk of branched crypts are unstained and can represent intermediates in either fusion or fission we propose the agnostic term FUFI (Fusion or Fission) to describe these transition forms.

### Crypt fusion and fission are regulated independently

To calculate crypt fusion rates heterotypic and homotypic FUFI forms were first evaluated for STAG2 and KDM6A loss by scoring around 3.9 ×10^6^ and 1.8 ×10^6^ crypts from 53 and 102 individuals each, respectively. In total, over 28,000 FUFIs were evaluated. This identified 151 FUFIs with STAG2 loss of 18,928 clones analysed and 63 FUFIs with KDM6A loss of 5,353 clones analysed (Figure 3D). These could be found as single events or as part of multicrypt clones (Figure 3E).

Importantly, while all M/W FUFIs are considered fusions and are directly observable, M/M FUFIs can be either fissions or fusions. The fusion rate is potentially accessible if the relative contribution of M/M FUFI events to the total number of fusions (termed *k*) can be determined. The value for *k* is calculable on the basis that fusions are classed as either M/M or M/W depending on the number of M and W neighbours present at the onset of fusion. To obtain this data, M/M and M/W FUFI clones were identified within multicrypt clones where the status of all their neighbours could be scored as W or M (totalling 4, 121 and 49 for mPAS, STAG2 and KDM6A, respectively) (Figure 3 F&G). Single branched crypts, containing clones in either branch, that possessed only W neighbours were also scored. This analysis revealed *k* to be 0.03 for mPAS. Within the smaller clones that are characteristic of neutral clonal marks such as mPAS, most fusion events are W/M leading to low values of k. Biased clonal marks generating large expansions will have a greater contribution from M/M fusions. Correspondingly the *k* values for STAG2 and KDM6A are 0.26 and 0.35, respectively (Figure 3H). Rates of fusion using these values of *k* can be estimated by: (i) assuming equal duration (i.e. the time window during which FUFIs are detectable as intermediate forms) for both fusion and fission; (ii) using the fission rate that is independently derived from changes in patch size distributions with age (see methods).

This analysis indicated similar crypt fusion rates for mPAS, STAG2 and KDM6A of 0.3% per year (95% CI: 0.1-0.6), 0.4% (95% CI: 0.3–0.7) and 0.7% (95% CI 0.3-1.4), respectively that were not significantly different (Figure 3I). Comparison of mPAS fission and fusion rates, (the former is 0.7% per year; 95% CI: 0.5-0.9) show that these closely correspond. This suggests that for neutral mutations the rates of the two processes are balanced and will act together to maintain constant crypt numbers across the tissue as has been suggested previously (22). However, for biased mutations that, by definition, show elevated fission rates there appears to be no evidence for a compensatory elevation of the fusion rate. These analyses suggest that fission and fusion are independent processes and not coordinately regulated.

### Crypt diffusion accommodates new crypts throughout life

A striking feature of larger patches is that mutant crypts have over decades populated the territory initially occupied by multiple crypts without a significant increase in crypt density. In the absence of appreciable merging of crypts by fusion this suggests local adjustments or remodelling to allow crypts to be displaced from the growing focus and return to near ambient densities. With this rationale we considered the possibility of random crypt movement in the form of a diffusion process.

Translated into the colonic epithelium the density of crypts in the ‘space’ of the surrounding stroma is the material being diffused. Each mutant clone comprises a source and new crypts created by successive fission events create a perturbation in local crypt density. Crypt fission thus produces a local decrease in crypt spacing which is relieved by outward diffusion, notably of both the mutant and surrounding wild type crypts, until a new balance is reached (Figure 4A). The system dependent diffusion coefficient (change in area per unit time) can be estimated based on measured crypt spacing and theoretical bounds on patch age.

**Figure 4.**
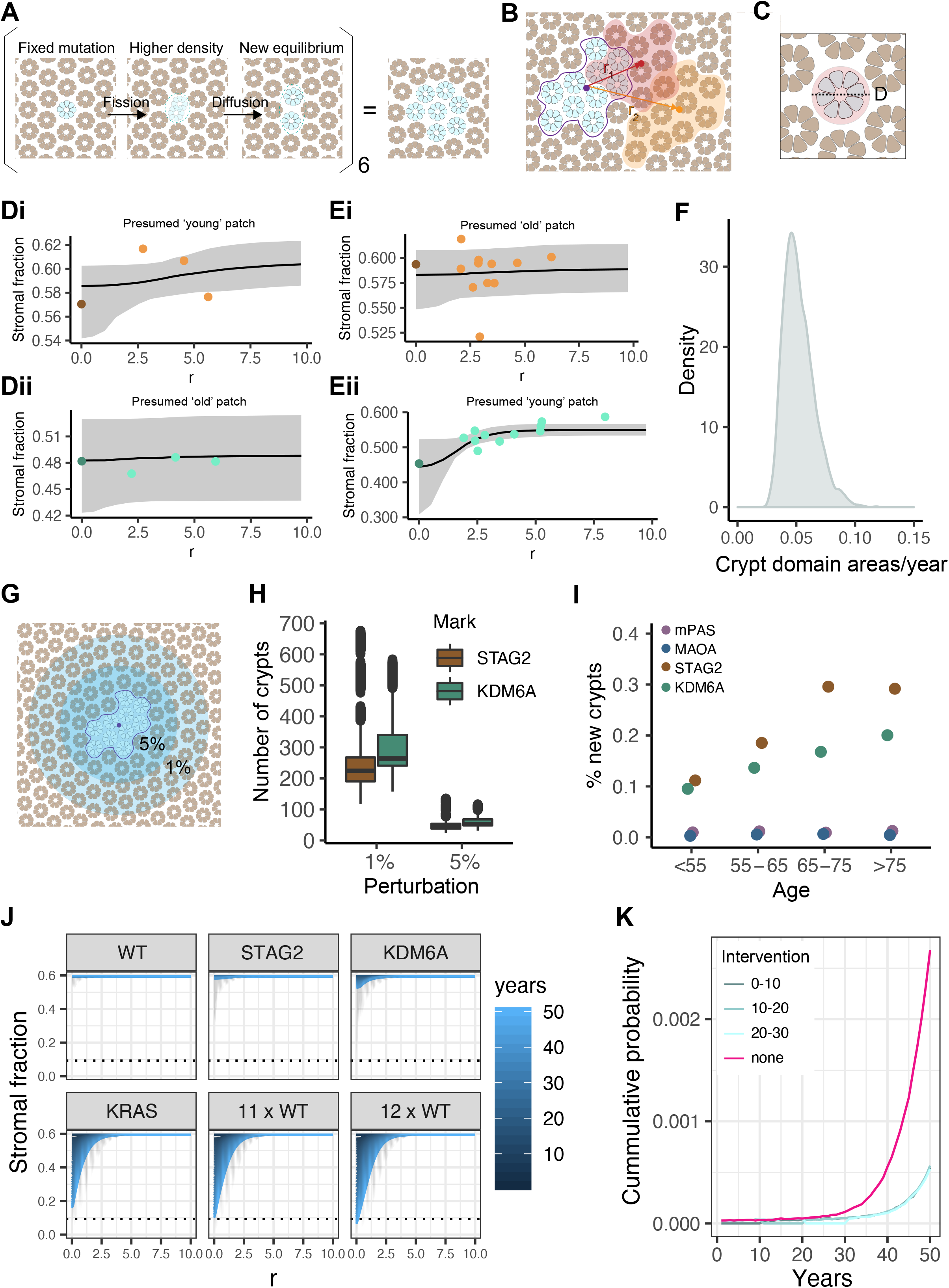
Evidence for a crypt diffusion process. (A) Schematic representation of the proposed crypt diffusion process. Fission of a clone results in higher local density which is then relieved by diffusion. (B) Schematic representation of areas measured to assess packing of mutant patch and surrounding crypts in rolling windows placed at different distances (R) from centroid of mutant patch. (C) Schematic showing crypt cross section with area of surrounding stroma that defines a crypt domain. (D) Examples of fitted diffusion process to achieve ambient density when comparing patches of 10 crypts when moving from mutant clone to three adjacent control groupings. (i) STAG2, (ii) KDM6A. Points are data derived, line is best fit, grey shading is 95% CI: presumed “young” patch and bottom: presumed “old” patch for i) STAG2 and ii) KDM6A. (E) As (D) but with rolling window data. (F) Density plot of values obtained for the diffusion coefficient in human colonic epithelium. (G) Schematic representation of area of crypts affected if space is decreased by 5% or 1% surrounding a mutant patch of 10. (Crypt numbers not to scale). (H) Boxplot showing the simulated numbers of crypts affected if spacing is decreased by 1% or 5% respectively due to addition of a patch of 10 mutant crypts. (I) Dotplot showing median frequency of newly generated crypts for mPAS, MAOA, STAG2 and KDM6A across four age bins. (J) Line graphs showing the stromal fraction resulting from crypt diffusion and different crypt fission rates corresponding to the homeostatic (wild-type) rate as well as those associated with STAG2, KDM6A and KRAS and multiples of 11 and 12 of the WT rate. Dotted line = whitespace fraction calculated from optimal hexagonal packing of circles. R = distance from centroid of patch in crypt domains. (K) Line graph showing results from simulations to calculate the cumulative probability of a clone growing into a lesion over time. Simulations were performed using the fission rate associated with KRAS activation (none) and are shown alongside the results obtained for simulation inclulding therapeutic crypt fission inhibition applied for a decade either immediately after mutation acquisition (0-10), after 10 years (10-20) or 20 years (20-30).

To find evidence supporting a diffusion type process analysis was focused on clones comprising ten crypts, the largest size for which sufficient data could be obtained for STAG2 and KDM6A (24 and 20 patches, respectively). For each mutant clone, and for three arbitrary control groups of ten adjacent wildtype crypts, each crypt was spatially mapped in X/Y coordinates. Areas of individual crypts as well as total patch area were measured and the distance between mutant clones and wild type patch centroids was determined. Finally crypt domains were defined as the area occupied by both crypt and surrounding stroma (Figure 4B & C).

The age of each clone was inferred in tandem with the diffusion coefficient and the ambient stromal fraction for each slide to define an overall tissue process that best fits the data (Supplemental Figures 4 & 5). The tissue-intrinsic diffusion coefficient describes how the tissue responds to the mutant clone by dispersing the burden of decreased stromal fraction into the surrounding tissue over time. For older clones we expect the system to be close to the ambient density while for younger clones the perturbation to the local stromal fraction may still be evident. Examples of both presumptive young and old clones were readily detectable in our samples (Figure 4D and Supplemental Figures 6 & 7).

For a subset of 7 patches (4 for KDM6A, 3 for STAG2) a more detailed rolling window analysis was performed in which the above approach was applied where fields of ten crypts were moved outwards from the mutant clone. Again evidence of a reduction in stromal fraction consistent with perturbation in younger clones was observed (Figure 4E and Supplemental Figures 6 & 7). The diffusion coefficient was found to be 0.055 crypt domain areas/year (95% CI 0.053 – 0.057) (Figure 4F). Model comparison were performed using leave-one-out cross-validation: exhaustive resampling of test-train perturbations was approximated using the full Markov chain Monte Carlo output to estimate pointwise out-of-sample prediction accuracy. Reassuringly this yielded a significantly worse fit than the experimental comparisons (Supplemental Figure 8).

This inferred diffusion process can be used to define the number of crypt domains affected to accommodate a new clonal expansion. The denser the new equilibrium of packing, the fewer crypts are perturbed to accommodate a clone of given size (Figure 4G & H). For example, the model suggests that patches of ten KDM6A^−^ crypts would require 264 crypt domains to undergo a 1% reduction in their spacing, while a 5% reduction would only require 52 crypt domains (Figure 4H).

### Defining a homeostatic threshold

Limited evidence suggests that there are no significant age-related changes in colon length and crypt density (20). With respect just to STAG2 and KDM6A mutations the relatively few new crypts arising during life could be easily accommodated by crypt movement. By the time individuals exceed 75 years of age, for every 10^5^ crypts, fission has added only approximately 200 and 290 new STAG2^−^ and KDM6A^−^ crypts, respectively (Figure 4I). However, it seems highly probable that additional genetic variants will also promote fission to different degrees. The potential for diffusion to locally balance this process as fission rate increases was investigated.

Simulations were performed escalating the homeostatic fission rate of 0.7% per year. When fission rates remain below approximately 11-fold above homeostasis, diffusion can generate enough space to accommodate newly generated crypts (Figure 4J). Higher fission rates result in a proportion of clones reaching a threshold of maximum packing density within which crypts are directly touching. This suggests a potential boundary for polyp growth, that is dependent on the physical processes of fission and diffusion. Interestingly, *KRAS* activating mutations, which we previously found to accelerate fission around 10-fold (1), mostly fall just below the threshold calculated here (Figure 4J and Supplemental Figure 9). Nevertheless, our model predicts each KRAS-mutant clone to harbour a cumulative probability of 0.25% of growing into a visible lesion 50 years after mutation acquisition. A therapeutic intervention inhibiting crypt fission for 10 years could reduce this to approximately 0.05% (Figure 4K).

## Discussion

Several studies have shown that advantaged mutations can generate large epithelial expansions (17). In the normal adult colon this is mediated by crypt fission. Perhaps because this is an infrequent event there has been little consideration of the compensatory mechanisms that allow newly made crypts to be accommodated within a fully formed epithelium. However, here and previously it has been shown that biased mutations can significantly increase crypt fission rates to generate large crypt expansions that are appropriately distributed within the tissue (1). In contrast increased glandular fission rates are also known to drive the overgrowth of adenomas and CRCs suggesting that differences in the rate of fission or the response to it must differ between normal and neoplastic tissues (12–15).

In considering the epithelial responses that compensate for elevated fission rates we first validated a new clonal mark, based on loss of KDM6A, that displayed advantage in the process. This together with STAG2 provided two gene specific assays with 5- and 3-fold increase in homeostatic fission rates. Comparing the configuration of the size and frequency of clones for the two genes demonstrates the different strategies by which age-related mutational burden can be achieved. STAG2 has the higher mutation rate and generates many relatively small clones while KDM6A generates fewer but larger expansions. A corollary of the exponential growth of patches as their size increases is that larger patches will tend to be the most recent and therefore most likely to contain evidence of local adaptation to accommodate new crypts.

The recently recognised process of crypt fusion offers a potential mechanism to compensate for fission events (21). Assuming they had equal rates in homeostasis they could effectively balance crypt numbers on a population basis (22). In considering fusion as a mechanism to accommodate new crypts, a baseline estimate was first established here and found to approximate that for fission. However, no upregulation of fusion accompanying the local expansions resulting from STAG2 and KDM6A mutation was found. Conceivably other mutations may impact fusion to ease local packing but it does not appear necessary to do so.

Multicrypt clones that form over decades inevitably populate the territory previously occupied by multiple independent crypts. In seeking to understand this dispersal we looked for and found evidence of a diffusion-type process in a subset of clones. These are consistent with a recent clonal expansion ‘caught in the act’ of being restored to an ambient crypt density. The behaviour captured probably reflects a passive dispersal mechanism rather than actual diffusion and must be accompanied by some level of stromal turnover.

The diffusion coefficient defines the rate of movement of crypt domains and the size of the larger impacted zone that is required to absorb new crypts. Parameterising the process allows testing of the robustness of the tissue to deal with localised accelerated growth conferred by biased mutations such as STAG2 and KDM6A. From this analysis the homeostatic dispersal mechanism seems over engineered and able to accommodate increased fission rates of more than 10-fold above baseline. Even for mutations that generate higher crypt fission rates only the fastest growing clones would overgrow the available space. For example, around 5% of clones carrying a gene mutation that confers a 19-fold increase in fission rate would reach a threshold where they lack a stromal domain between crypts and overgrow the available space.

The actual threshold at which recent clonal expansions become recognisable as pathologies may be lower than the extreme one applied here. However, the implication remains that the distinction between phenotypically normal clones and those forming overt pathologies may be determined solely by a probabilistic process in which a recent succession of fission events overcome homeostatic dispersal mechanisms.

Activating mutations of KRAS have been described in normal colonic epithelium. Using targeted sequencing we previously identified high mutant allele frequencies for KRAS G12 and G13 mutations in a proportion of patients and inferred that a 10-fold increase in fission rate was required to achieve them. It is intriguing that activating mutations of KRAS come close to achieving the fission rates capable of reaching the extreme threshold defined here. If the actual threshold is lower, or if different KRAS mutations confer different fission rates, then a proportion of such clones will overgrow. Of note, KRAS is commonly mutated in a broad spectrum of benign and premalignant pathologies such as serrated lesions and adenomas that may arise at least in part due to this dispersal threshold being reached (30–34).

It is known that a proportion of adenomas spontaneously regress when observed in longitudinal studies (35–37). One plausible explanation for such phenomena is that these lesions first develop due to reaching the threshold resulting in local overcrowding but are transient because of ongoing crypt dispersal.

Loss of function mutations affecting the APC tumour suppressor gene are also initiated and expanded by glandular fission (12–15). Mutation of both APC and KRAS is frequently found in CRCs. The two pathways activated are known to interact at the molecular level (38). It is likely their combined activation will also synergise to further elevate gland fission rate and promote overgrowth as fully neoplastic CRCs develop.

Clonal expansions without an overt phenotype but predisposing to oncogenic transformation may explain the interpretation that colorectal cancers arise as a single expansion event when a combination of factors reaches a critical threshold (19). This ‘Big Bang’ model requires that most truncal mutations required for neoplastic transformation are acquired within a single cell. Accumulating such multiple hits will be facilitated probabilistically by the growth of large epithelial expansions that act to increase the population available for secondary mutation. However, the trigger point for transformative growth may depend on additional non-mutational stochastic processes of which achieving a growth rate that overruns local dispersal mechanisms is one.

Obesity, a known risk factor for CRC, is known to be accompanied by an increase in crypt fission rate (39). More specifically diets deficient for methyl donors are known to reduce crypt fission rates in the mouse (40). An implication of the colon having the capacity to absorb many more new crypts than are produced by homeostasis is that modest and time limited reductions in the fission rates resulting from pro-oncogenic mutations may not only slow the growth of lesions but prevent them from forming at all.

## Materials and Methods

### Human tissue

Normal colon tissue samples were obtained from both Addenbrooke’s Hospital Cambridge and Norfolk and Norwich University Hospital under full local research ethical committee approval (Approval document IDs 15/WA/0131 & 17/EE/0265 as well as 06/Q0108/307 & 08/H0304/85, respectively) according to UK Home Office regulations. A total of 228 individuals were included in the study with an age range of 8–93 years. Colectomy specimens were fixed in 10% neutral buffered formalin. From areas of tissue without macroscopically visible disease mucosal sheets were removed from the specimens and embedded *en face* in paraffin blocks. For standard stainings, sections were cut from each sample at 5 μm thickness and mounted onto charged glass slides.

### mPAS staining

Sections were dewaxed and rehydrated on a Leica Multistainer ST5020 before washing in 0.1 M Acetate buffer pH 5.5 at 4 °C for 5 minutes. This was followed by oxidation in 1 mM sodium periodate buffer at 4 °C for 10 minutes before washing in 1% glycerol for 5 minutes. Then, three washes were performed in ultra-pure water for 5 minutes in total, followed by staining in Schiff’s reagent for 15 minutes. Sections were then washed again in ultra-pure water and counter-stained in Mayer’s Haematoxylin for 40 seconds. After another wash in ultra-pure water and brief blueing in tap water and a final rinse in ultra-pure water was performed. This was followed by dehydration, clearing and mounting in DPX on the same Leica Multistainer.

### Immunohistochemistry

#### Standard protocol

Sections were dewaxed and rehydrated on a Leica Multistainer ST5020. This was followed by heat induced epitope retrieval in citrate buffer (10 mM sodium citrate, pH6) in a medical pressure cooker and washing in PBS. All subsequent washes were 3 x 5 min in PBS-T (PBS with 0.05% Tween-20). After a 15 min incubation with 3% H_2_O_2_ in methanol and a wash, slides were incubated with blocking buffer (PBS-T with 10% Donkey Serum, Dako) for 30 min. This and all following incubations were performed at room temperature in a humidified slide box. Slides were then incubated with the primary antibody (see table 1 in appendix) for one hour at room temperature or overnight at 4 °C. After a wash, slides were incubated with the secondary antibody (biotin-SP-conjugated AffiniPure donkey anti-mouse, anti-rabbit or anti-goat, Jackson ImmunoResearch, all 1:500 in PBS-T) for 40 min. Following a wash, slides were incubated with Vectastain® Elite® ABC reagent (Vector Laboratories) for 40 min.

This was followed by a final wash and immunoperoxidase detection using a liquid DAB + substrate chromogen system (Dako). Finally, sections were counterstained with Mayer’s Haematoxylin, dehydrated and mounted in DPX on the Leica Multistainer.

#### Adaptations for laser capture slides

For sections for laser capture microdissection (LCM) the above protocol was modified as follows: Tissue was cut at 10 μm thickness onto UV-irradiated PEN membrane slides (ZEISS). Heat induced epitope retrieval was performed in citrate buffer in a water bath at 76 °C for 16 hours. Counterstaining with Mayer’s Haematoxylin was performed manually for 15 seconds followed by blueing in tap water for 1 minute and drying at room temperature. Slides were stored at room temperature (short-term) or −20 °C (long-term).

### LCM and sequencing of KDM6A-negative patches

#### LCM and DNA extraction

On stained FFPE human colonic tissue sections, crypts were harvested into lids of 0.2 mm radius PCR tubes using a Leica LMD7000 Laser Microdissection System. 10 μl of Proteinase K solution from the Arcturus® PicoPure® DNA Extraction Kit (ThermoFisher) were added and tubes centrifuged in a mini-centrifuge for PCR tubes. Following lysis for 3h at 65 °C in a standard PCR block the enzyme was inactivated by incubation at 95 °C for 10 min. Samples were kept at 4 °C until PCR.

#### Assessment of DNA quality

For assessment of optimal amplicon size, DNA extracted from FFPE sections as described above was diluted to equivalents of 5, 20 and 100 crypts and used for PCR reactions containing primers at 1 μM concentration (see table 2 in appendix), 0.5 mM dNTPs (New England BioLabs), 5 μl 5X Phusion® HF Reaction Buffer (New England BioLabs), 1 U of Phusion® High-Fidelity DNA Polymerase (New England BioLabs) and nuclease-free H_2_O (Ambion) to make up a volume of 25 μl. PCR cycling was performed at 95 °C for 2 min for one cycle, followed by 35 cycles at 95 °C for 10 s, 60 °C for 10 s and 72 °C for 15 s. The final cycle was followed by a 5 min extension at 72°C.

#### Pre-amplification

Due to low quantities of input DNA a pre-amplification PCR was performed. All primers were designed using Primer3 (41). They included Fluidigm CS1 and CS2 tags and were ordered from Sigma (see appendix table 3). Reactions were performed in multiplex primer groups with primers each at 1 μM concentration (group 1: primers 1,4,7…group 2: 2,5,8… group3: 3,6,9…). The LCM DNA sample was diluted such that each PCR reaction contained 10 μl DNA sample. In addition, each reaction contained 0.5 mM dNTPs (New England BioLabs), 5 μl 5X Phusion® HF Reaction Buffer (New England BioLabs), 1 U of Phusion® High-Fidelity DNA Polymerase (New England BioLabs) and nuclease-free H_2_O (Ambion) to make up a volume of 25 μl. PCR cycling was performed at 95 °C for 2 min for one cycle, followed by 35 cycles at 95 °C for 10 s, 60 °C for 10 s and 72 °C for 15 s. The final cycle was followed by a 5 min extension at 72°C. Samples were then treated with ExoSAP-IT enzyme (2 μl of enzyme for 5 μl of sample, ThermoFisher) at 37 °C for 15 min followed by a 15 min inactivation at 80 °C. Samples were then diluted 1:10 in DNA Suspension Buffer (Teknova) and stored at 4 °C until further processing.

To obtain adequate amounts of DNA for sequencing, the pre-amplified products were further amplified using the Fluidigm Access Array™ according to the supplier’s protocol

#### Barcoding and Next-generation sequencing

A unique Fluidigm barcode was added to each sample by PCR in 10 μl reactions using the Fast Start High Fidelity PCR System (Roche) containing: 400 nM barcoding primers, 1 μl diluted PCR product, 1X Fast Start HF buffer without MgCl_2_, 4.5 mM MgCl2, 5% DMSO, 0.2 mM dNTPs each, 0.05 U/μl High fidelity enzyme mix and nuclease-free H_2_O (Ambion) to make up a reaction volume of 10 μl. PCR cycling was performed using the protocol: 95°C for 10 min followed by 15 cycles at 95 °C for 15 s, 60 °C for 30 s and 72 °C for 60 s. The final cycle was followed by a 3 min extension at 72 °C. Samples were then pooled, purified using a Clean & Concentrator Kit (Zymo Research) and primer dimers eliminated by broad range (200-400 bp) size selection using a PippinBlue (Sage Science). Samples from one Fluidigm Access Array™ were sequenced on one lane using 150 bp paired-end sequencing with 10% PhiX in-house on the Illumina MiSeq platform.

#### KDM6A mutation calling

Fastq files were converted to .BAM files and analysed using a PERL script (available upon request) by Dr Richard Kemp. Briefly, from the bulk of reads the script first identifies reads corresponding to the amplicons of interest by pulling out reads starting and finishing with the expected sequence as well as containing an expected stretch of sequence in the middle. For these reads, at every nucleotide position outside of the primer sequence, the number of reads corresponding to the reference genome as well as the number of reads containing a base change at that position are recorded. This enables calculation of the noise at every position. Candidate mutations were identified when the mutant allele frequency was either >4x the mean of the noise at that position or >3.29 x the standard deviation at that position (p≦ 0.001). True mutations were called if they were present in all samples originating from the same patch in serial sections but absent in all wild-type samples from the same sections. This analysis was complemented by manual inspection of reads using the Integrative Genomics Viewer by the Broad Institute (42).

### KDM6A clone data acquisition

#### Individual clones and patches

WPC and PPC numbers as well as multicrypt patch sizes were manually scored in stained tissue sections using a standard brightfield microscope. Sections were scanned at 20X using a Leica Aperio AT2 scanner and stored as .svs files on a CRUK Cambridge Institute server. Scans were examined online using the Aperio eSlide Manager (Leica) or downloaded and annotated using QuPath (43). To count the total number of crypts on each section, .svs files were analysed with the DeCryptICS neural network developed by Dr Edward Morrissey and Dr Doran Khamis (https://github.com/MorrisseyLab/DeCryptICS, manuscript in preparation).

#### Quality control

Within the dataset two individuals aged 37 years with extreme average patch sizes were identified as outliers with respect to that measure and not included in subsequent analyses of patch sizes, fusion rates and newly generated crypts.

#### Scoring crypt fusion

To score crypt fusion, the term ‘FUFI’ was invented to agnostically refer to structures that could represent a crypt undergoing fusion or fission in *en face* human colonic tissue sections. A FUFI was defined as two crypts joined together without any visible gap but with two clearly distinguishable lumina. For stained sections, FUFIs can consist of two wild type crypts (WT FUFI), two mutant crypts (mut FUFI) or a mutant and wild type crypt (M/W FUFI). To count the total number of FUFIs on each section, .svs files were analysed with DeCryptICS followed by manual annotation in QuPath to classify FUFIs as WT, M/M or M/W.

#### Measurement of crypt packing and stromal fraction

Crypt domains were defined as a crypt and its surrounding stroma. Crypt domains were drawn onto scans of tissue sections (.svs files) using QuPath software, whereby lines were placed onto the stroma centrally between neighbouring crypts. Initially, to measure the stromal fraction in STAG2- and KDM6A-mutant patches comprising 10 crypts (24 and 20 patches, respectively), the total area of crypts was divided by the total area of crypt domains within the patch. The same measurements were also performed for adjacent groups of 10 wild type crypts, totalling three per mutant patch.

To obtain measurements for inference of the diffusion coefficient, crypt areas, x- and y-coordinates, total area of crypt domains as well as the distance to the centroid of the mutant patch, r, were obtained for the mutant as well as surrounding wild type patches. In addition, for 7 patches (3 STAG2, 4 KDM6A) the same analysis was also performed as a rolling window, moving outwards from the mutant patch. This included, for each mutant patch, surrounding two areas each comprising 3 mutant and 7 WT, 2 mutant and 8 WT and 1 mutant and 9 WT crypts as well as five surrounding WT patches at varying distances (combined: 1 data point for the mutant patch and 11 for surrounding patches).

### Mathematical inference of stem cell dynamics and crypt fission

The stem cell dynamics and fission rate associated with loss of KDM6A were mathematically modelled as previously described (1). This includes modelling of mutation acquisition as well as competition with other stem cells in the crypt, which can result in clone fixation. Subsequent crypt fission was modelled as a Yule-Furry pure birth process.

### Mathematical modelling of crypt fusion

See mathematical appendix.

### Mathametical modelling of crypt diffusion

See mathematical appendix.

## Acknowledgements

The authors thank the Biobanking, Histology and Genomics cores at the CRUK Cambridge Institute for technical support. The authors acknowledge the contribution and support to this project provided by the Norwich Research Park BioRepository (Human Tissue Authority licence number 11208; NRES REC nos 06/Q0108/307 & 08/H0304/85+5), a facility supported by the BBSRC, the Norfolk and Norwich University Hospitals NHS Foundation Trust and the University of East Anglia. Equally, the authors acknowledge contribution and support provided by the Addenbrooke’s Human Research Tissue Bank (NRES REC nos 15/WA/0131 & 17/EE/0265) that is supported by the NIHR Cambridge Biomedical Research Centre. Funding for this project was provided by a Wellcome Trust Grant (103805), Cancer Research UK, a Wellcome Trust 4-year Ph.D. studentship and a Medical Research Council Computational Biology Fellowship (MC_UU_12025).

## Notes

### Competing Interest Statement

The authors have declared no competing interest.

