## Supplemental materials for "A diffusion-like process enables expansion of advantaged gene mutations in human colonic epithelium"

#### Supplementary figure legends

**Supplementary Figure 1: Laser capture of KDM6A-negative patches**

(A) Image of gel electrophoresis of PCR products on FFPE DNA diluted to equivalents of varying amounts of crypts. (B) Image of IHC for KDM6A on laser capture slide (i) before and (ii) after laser capture microdissection. (C) Box plot showing reads obtained for each of the 24 amplicons used for sequencing of KDM6A-negative and control patches.

**Supplementary Figure 2: Size measurement of $STAG2^-$ and $KDM6A^-$ crypts**

Crypt areas in multicrypt clones containing 10 crypts that are $STAG2^-$ or $KDM6A^-$ negative (24 and 20, respectively). The area of individual crypts was determined using QuPath software. As controls, adjacent or nearby wild type groups of 10 crypts were analysed in the same way. Jitter plots show areas of crypts. Black line = median, **** = p-value = < 0.001.

**Supplementary Figure 3: Defining the FUFI**

(A) Representative image of KDM6A-positive bifurcating crypts sectioned longitudinally. (B) A FUFI is defined as two adjoined crypts viewed in a transverse section with two clearly discernible lumina lacking any separating gap.

**Supplementary Figure 4: Inferred ages of $STAG2^-$ clones**

Outcome of inference for the age of STAG2- clones in years covering the period from single crypt to a clone comprising 10 crypts. The theoretical density of a patch age arising from fission as a stochastic birth process (blue) is compared to ages inferred by constraining to the range [5,100] and using Bayesian priors peaked at half the patient age.

**Supplementary Figure 5: Inferred ages of $KDM6A^-$ clones**

Outcome of inference for the age of KDM6A- clones in years covering the period from single crypt to a clone comprising 10 crypts. The theoretical density of a patch age arising from fission as a stochastic birth process (blue) is compared to ages inferred by constraining to the range [5,100] and using Bayesian priors peaked at half the patient age.

**Supplementary Figure 6: Diffusion model prediction of stromal fraction changes for STAG2$^{-}$ clones**

Plots show the stromal fraction measured for STAG2- as well as surrounding wild type patches of 10 crypts. The radial distance r (in crypt domains) is measured from adjacent patch centroid to mutant patch centroid. The black line and grey ribbon is the median and 95% CI theoretical stromal fraction as fitted from the diffusion model. Green box: patches for which a “rolling window” was applied. For each mutant patch, surrounding measurements included two areas comprising 3 mutant and 7 WT, 2 mutant and 8 WT and 1 mutant and 9 WT crypts as well as five surrounding WT patches at varying distances (combined: 1 data point for the mutant patch and 11 for surrounding patches).

**Supplementary Figure 7: Diffusion model prediction of stromal fraction changes for KDM6A$^{-}$ clones**

Plots show the stromal fraction measured for KDM6A- as well as surrounding wild type patches of 10 crypts. The radial distance r (in crypt domains) is measured from adjacent patch centroid to mutant patch centroid. The black line and grey ribbon is the median and 95% CI theoretical stromal fraction as fitted from the diffusion model. Green box: patches for which a “rolling window” was applied. For each mutant patch, surrounding measurements included two areas comprising 3 mutant and 7 WT, 2 mutant and 8 WT and 1 mutant and 9 WT crypts as well as five surrounding WT patches at varying distances (combined: 1 data point for the mutant patch and 11 for surrounding patches).

**Supplementary Figure 8: Simulation of WT patch as source of new crypts**

Results for model fit comparison where the identity of the source patch (the patch undergoing clonal expansion) in each neighbourhood was shuffled randomly from the mutant patch to one of the adjacent patches. The fit, visualised by the kernel of the log posterior, for ten such shuffled data sets were and compared to the model fit for the original data (where the mutant patch is identified as the source of clonal expansion). The overlaid data table shows results from leave-one-out cross-validation to estimate the relative predictive accuracy of the original diffusion model compared to shuffled-data alternatives; negative elpd (expected log predictive density) for the shuffled data fits confirms a better predictive accuracy for the diffusion model that takes the mutant patch as the source of the clonal expansion.

**Supplementary Figure 9: Simulating the 'breaking point'**

Simulations were performed to find the fission rate at which clones may generate new crypts more quickly than can be accommodated by crypt diffusion. Line graphs show the stromal fraction resulting from crypt diffusion and different crypt fission rates in multiples of the homeostatic (wild type) rate (0.7% per year). Dotted line = stromal fraction calculated from optimal hexagonal packing of circles. r = distance from centroid of patch in crypt domains. Grey area = 95% CI.

#### Supplemental Figure 1

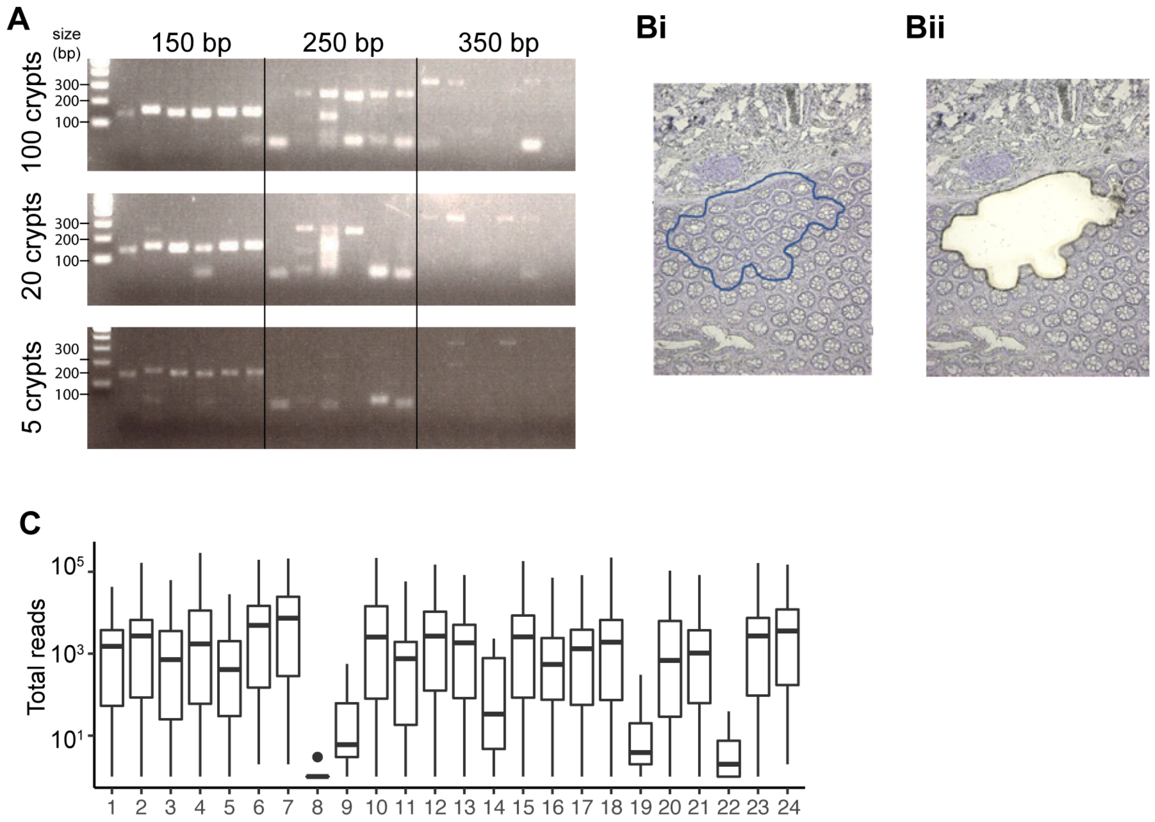

**Supplemental Figure 2**

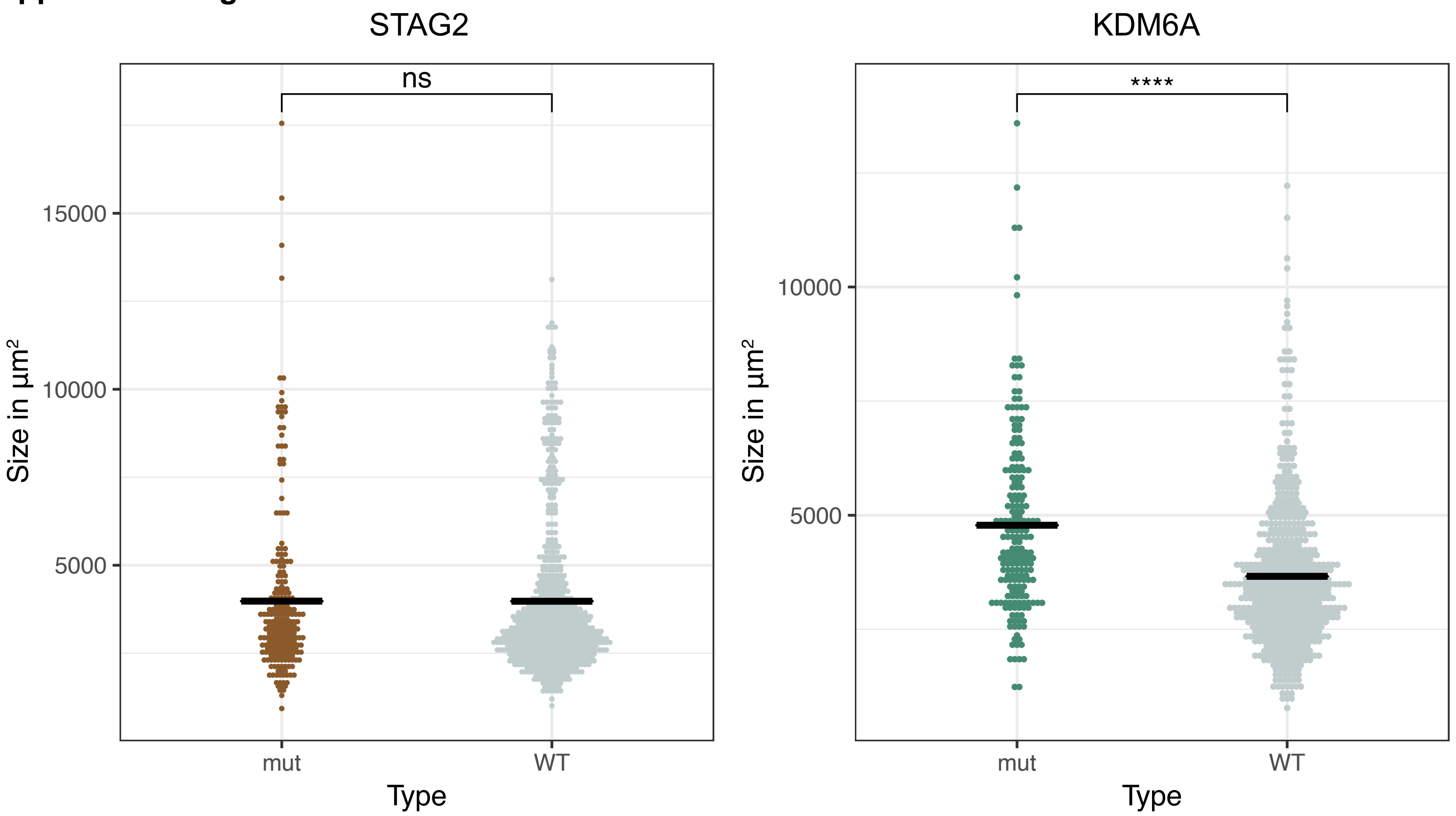

**Supplemental Figure 3**

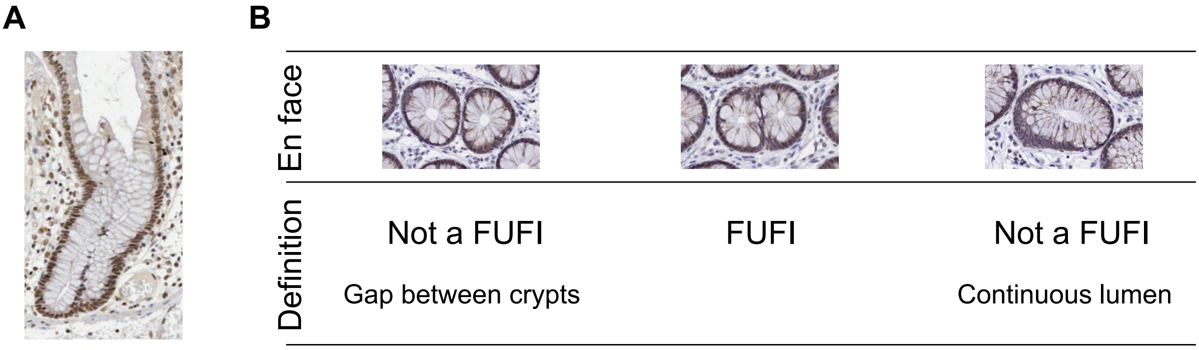

**Supplemental Figure 4**

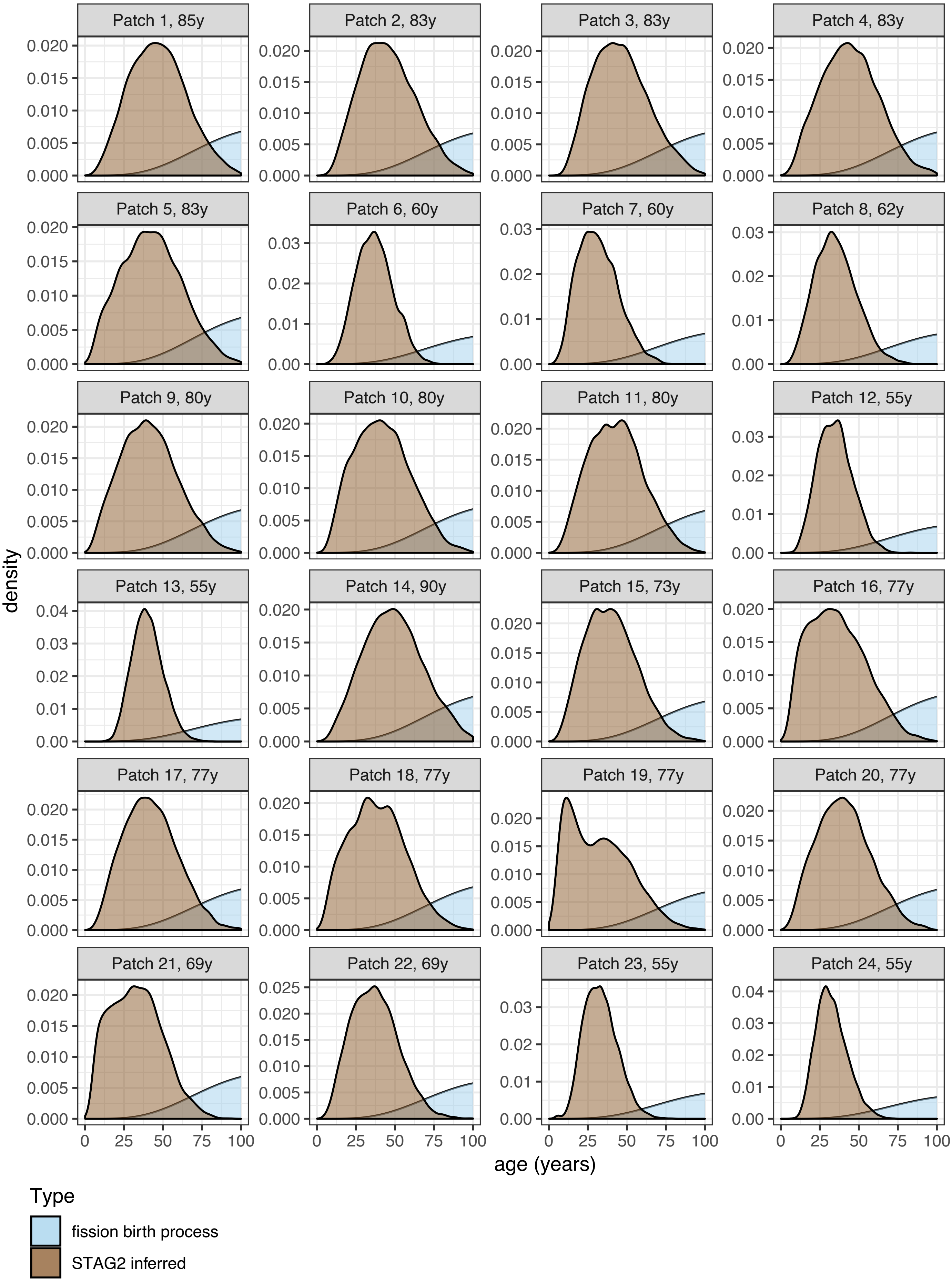

**Supplemental Figure 5**

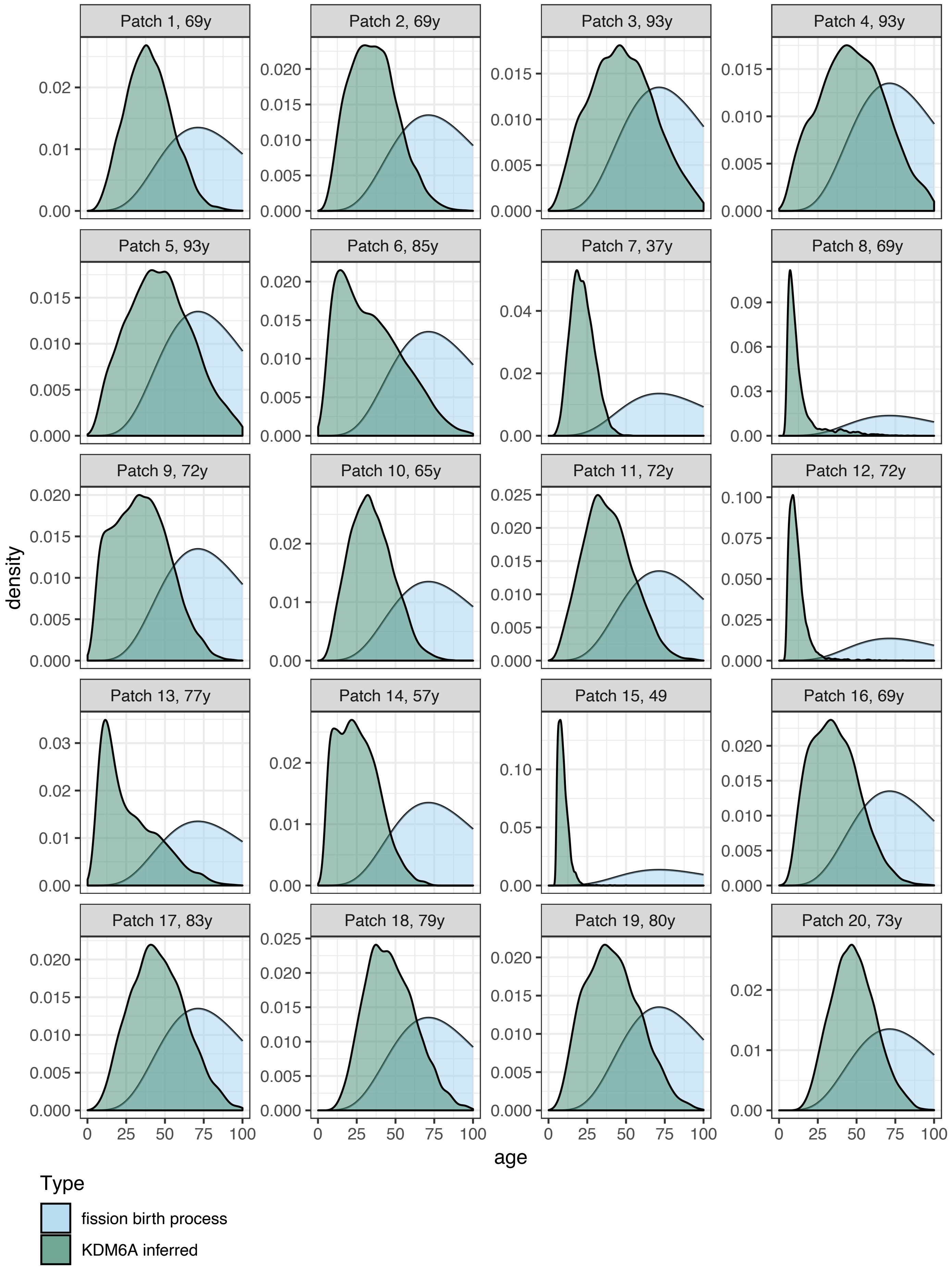

### Supplemental Figure 6

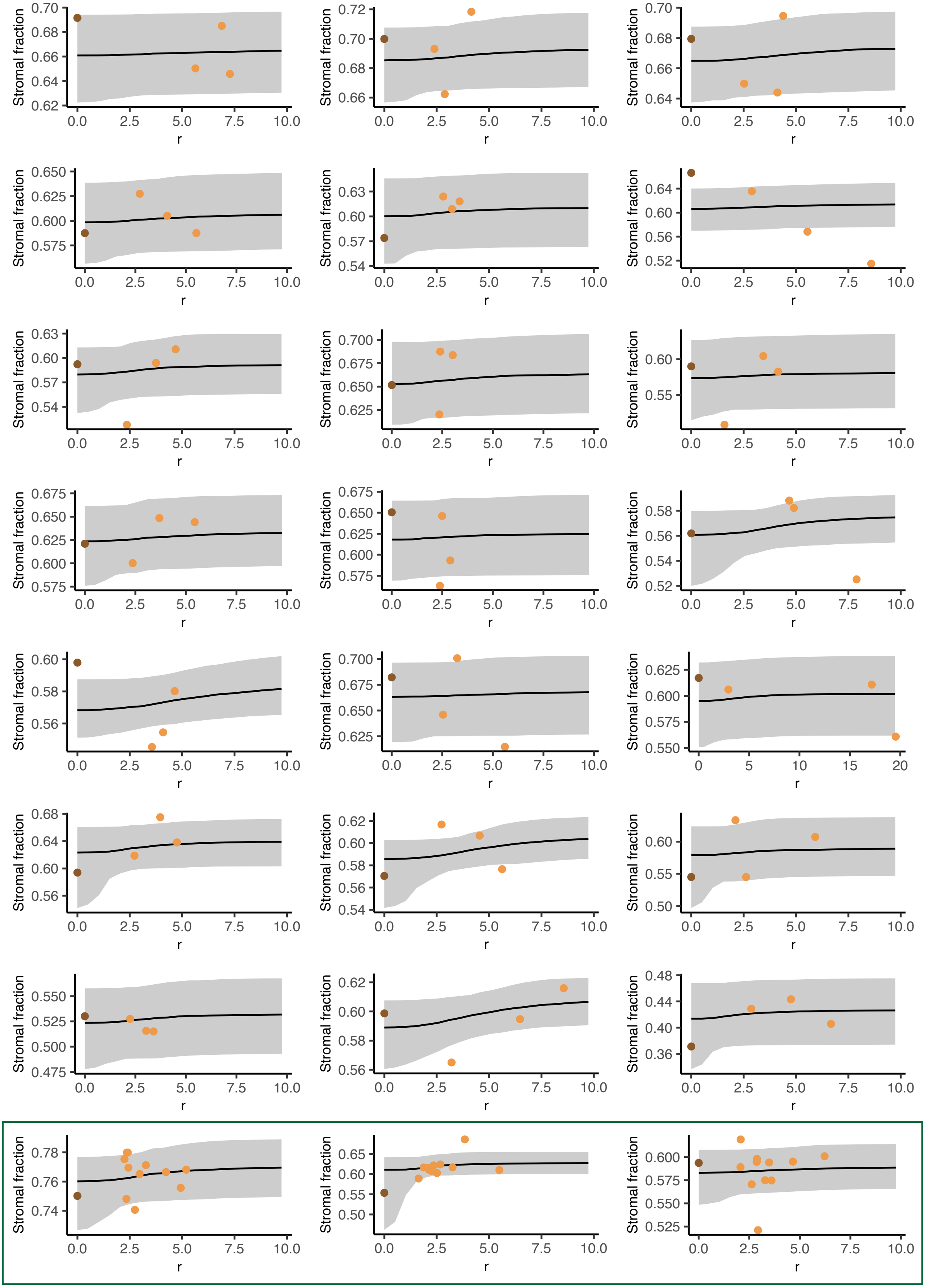

### Supplemental Figure 7

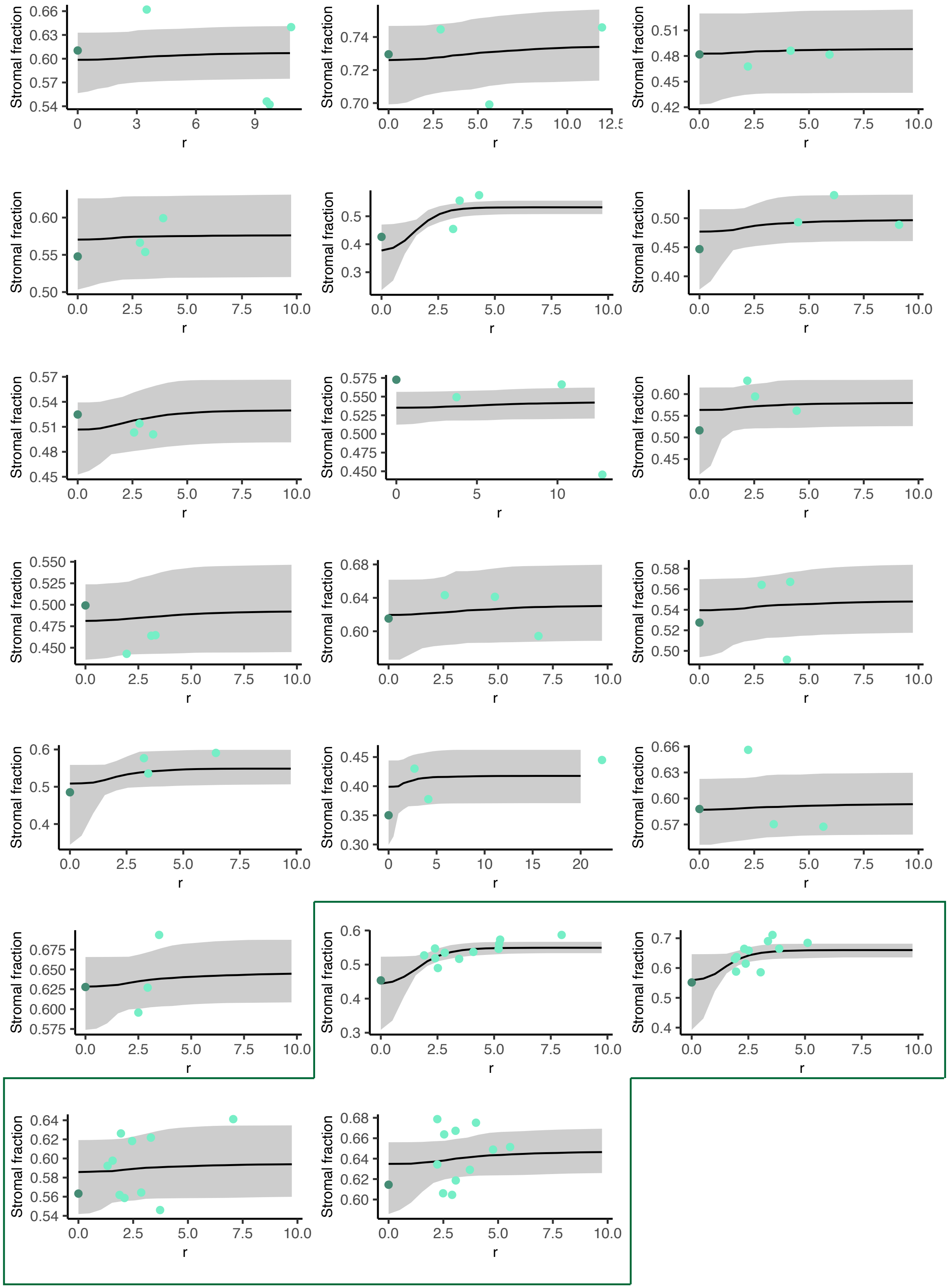

**Supplemental Figure 8**

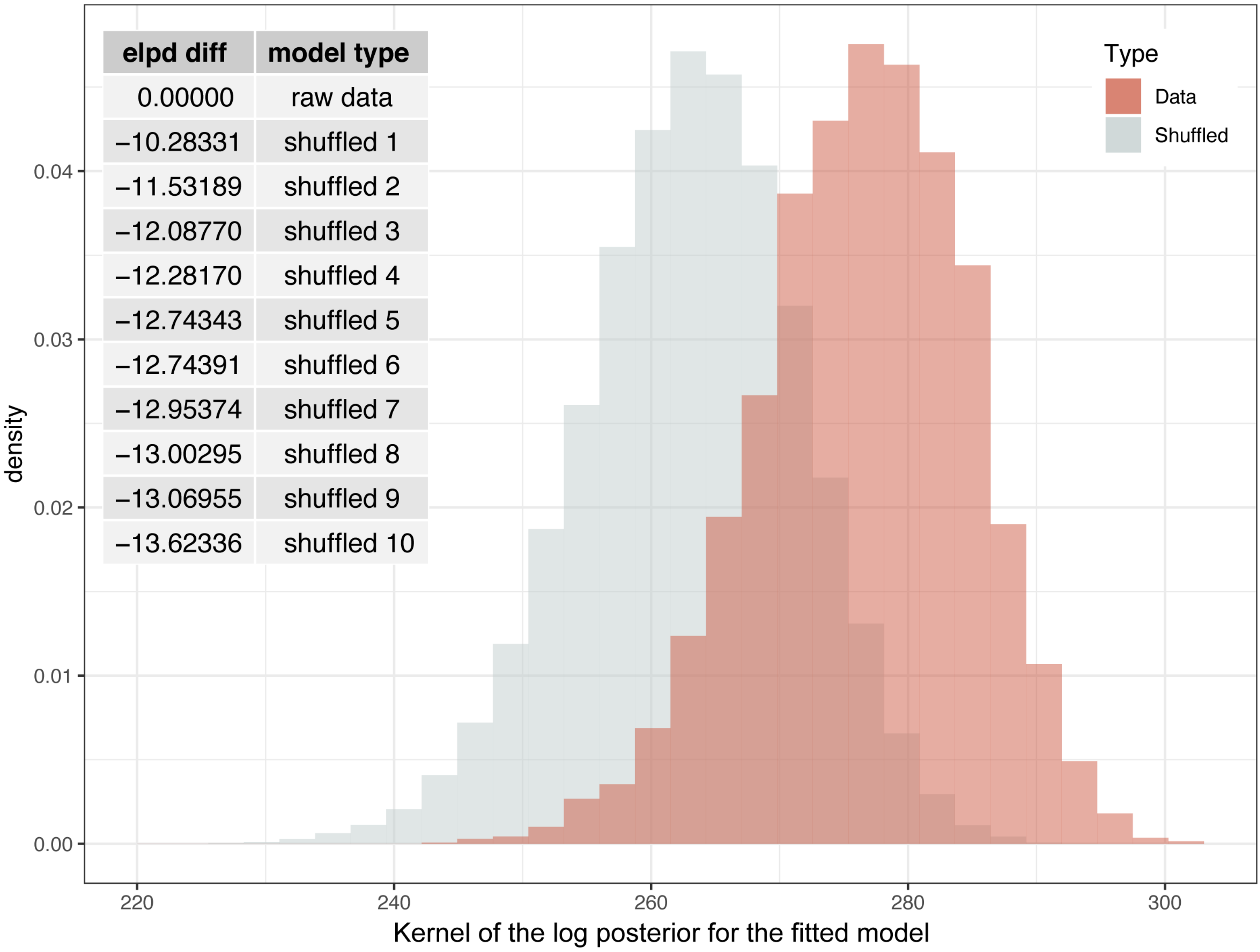

### Supplemental Figure 9

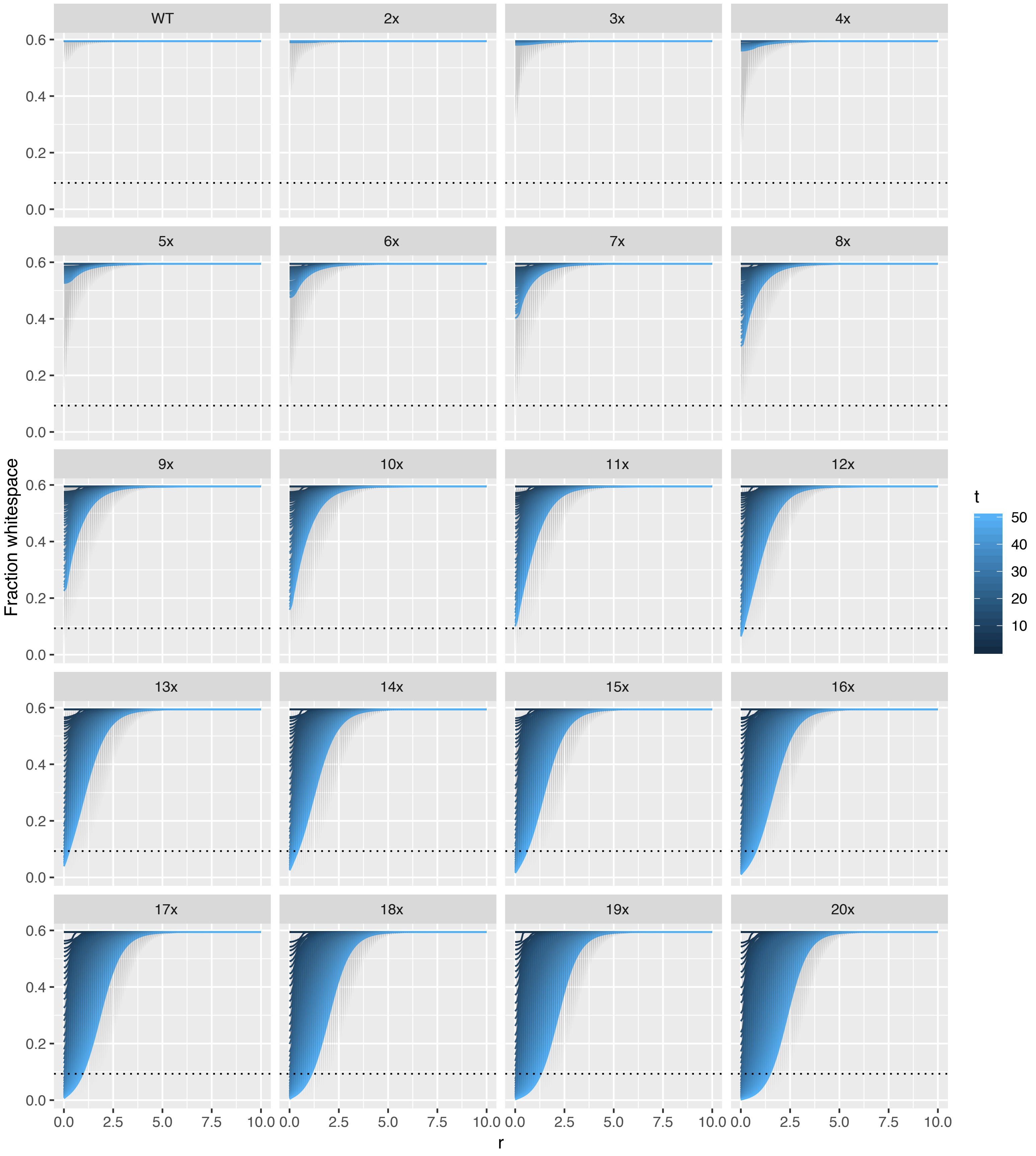

### Appendix

Table 1 **Summary of mutations identified in KDM6A$^{-}$ patches.**
*Ref = nucleotide in reference genome, freq = frequency, Alt = alternative nucleotide -> this is the mutation, Change = predicted consequence of the mutation. Frequencies derived from frequency of mutant reads in samples.*

| Gender | Sample | Exon | Ref | Alt | Alt freq | Maximum alt freq | Change |
|---|---|---|---|---|---|---|---|
| Male | 1A | intron 18 | A | G | 85.8 | 100 | Intronic |
| Male | 1B | intron 18 | A | G | 72.9 | 100 | Intronic |
| Male | 2A | intron 18 | A | G | 82.9 | 100 | intronic |
| Male | 2B | intron 18 | A | G | 85.3 | 100 | intronic |
| Male | 3A | 24 | C | A | 51 | 100 | S>STOP |
| Male | 3B | 24 | C | A | 0 | 100 | N/A |
| female | 4A | 25 | G | A | 15.8 | 50 | W>STOP |
| female | 4B | 25 | G | A | 10.4 | 50 | W>STOP |

Table 2 **Primary antibodies used for immunohistochemistry**

| Antigen | Antibody | Supplier | Titre |
|---|---|---|---|
| KDM6A | HPA002111 | HPA | 1:100 |
| KDM6A | #33510 | CST | 1:200 |
| MAOA | SC-271123 | Santa Cruz | 1:200 |
| STAG2 | LS-B11284 | LSBio | 1:1000 |

Table 3: **Primers used for amplification of FFPE to assess amplifiability.**

| Primer | Sequence (5' -> 3') |
|---|---|
| MAOA_150_F1 | ACCCATCAGTTACTCCTTCCC |
| MAOA_150_R1 | GGGATTAAAGCTGGGAGTTTCT |
| MAOA_150_F2 | TAGCAGGGCCTTGAATCTGT |
| MAOA_150_R2 | GATAGTGCCCAGAGTCACCA |
| STAG2_150_F1 | GGAGAAGAAGACACAGTTGGATG |
| STAG2_150_R1 | TTCTGTGAGGCATTTAGGGAAAA |
| STAG2_150_F2 | CCTATGCTCGCACAACTATGAG |
| STAG2_150_R2 | GGAAGCCACACATCCTCTCT |
| CASD1_150_F1 | ACCTGGAAACCCTATGCTCAA |

| | |
|---|---|
| CASD1_150_R1 | TGCAGCTATACATGCCAACC |
| LOC_150_F1 | TCGTCTGCTTCATCCTCCTC |
| LOC_150_R1 | GCCTAACATGCTTGGACCAC |
| MAOA_250_F1 | TGCAAGTCTTAGGTTGGTTGC |
| MAOA_250_R1 | TCAGTAATGGGTCATGTGCAAA |
| MAOA_250_F2 | AAGACATGTAGGGTTGGGGC |
| MAOA_250_R2 | CAGAACACCCTGCTCTAACCT |
| STAG2_250_F1 | GACTCTAAGGCCAGGTCAGG |
| STAG2_250_R1 | GGAGGTGAGTTGTGGTGTCT |
| STAG2_250_F2 | GCCTAATCATTCTCCCTGACCT |
| STAG2_250_R2 | TGGTGTCAAAATCCATTCCCTC |
| CASD1_250_F1 | GGTTAGAGGAAGACAAAAGTGGA |
| CASD1_250_R1 | CCTCAGTCCACACTTTGATACAC |
| LOC_250_F1 | AGCTTACCTCTTTGTCTCTTCCT |
| LOC_250_R1 | CAACCTCAAAGTATCACGTGGA |
| MAOA_350_F1 | TTCCTTCAGAAATTGAATCCTTG |
| MAOA_350_R1 | CCTGGGAGAAAGCAAAATCA |
| MAOA_350_F2 | TCCCGGAGTATCAGCAAAAG |
| MAOA_350_R2 | CATGAGAGACCCCCAAACAC |
| STAG2_350_F1 | TCCGAATATTTTTGGTGCATT |
| STAG2_350_R1 | CAGAGCCTTGATGAGTGCTG |
| STAG2_350_F2 | TCTGAAGGAATGCTATGGTATGAA |
| STAG2_350_R2 | TTGTCAAGGGTCATAGACACAA |
| CASD1_350_F1 | CTTTGGGAAGCTTTGCGTAAAA |
| CASD1_350_R1 | CGATTCAGGAAGATGTAAGCCA |
| LOC_350_F1 | TCAGGAATGATGGTCTACGTGA |
| LOC_350_R1 | TCTCAGCTCTATTCCGTGAGT |

Table 4 **Primers used for amplification of KDM6A.**
*Number = amplicon number, F = forward primer, R = reverse primer. Sequence includes Fluidigm CS adapters.*

| Primer | Sequence (5' -> 3') |
|---|---|
| 1_F | ACACTGACGACATGGTTCTACACGCTTTCGGTGATGAGGAAA |
| 1_R | TACGGTAGCAGAGACTTGGTCTCCGTACCTGTCCAGTCCG |
| 2_F | ACACTGACGACATGGTTCTACATCTTTCAGGGCAATTAAAGCATT |
| 2_R | TACGGTAGCAGAGACTTGGTCTACAACCTACCTTTAAACTAGACTCA |
| 3_F | ACACTGACGACATGGTTCTACAGTACAATTGGACCATGGCCA |
| 3_R | TACGGTAGCAGAGACTTGGTCTAGTGCAGAGGTATTACTACAACTT |
| 4_F | ACACTGACGACATGGTTCTACACAGGATGCCATTAAATGCTACTT |
| 4_R | TACGGTAGCAGAGACTTGGTCTTCTGGGGAAATATGTGGCTTT |
| 5_F | ACACTGACGACATGGTTCTACAATGCTGTGTCACATCCTCCA |

| | |
|---|---|
| 5_R | TACGGTAGCAGAGACTTGGTCTACTTGTTTGCTACCTCTACTCCT |
| 6_F | ACACTGACGACATGGTTCTACATGACAGATGAGACCAACAGGA |
| 6_R | TACGGTAGCAGAGACTTGGTCTCAGGCTGAGAGACGCTAGG |
| 7_F | ACACTGACGACATGGTTCTACACTGCCTACAAACTCAGTCTCTG |
| 7_R | TACGGTAGCAGAGACTTGGTCTCAGAAAAGGGTCCATTGGCC |
| 8_F | ACACTGACGACATGGTTCTACATAACCGCACAAACCTGACCA |
| 8_R | TACGGTAGCAGAGACTTGGTCTTCTCTCAAAGTGTATAAAACCCAGT |
| 9_F | ACACTGACGACATGGTTCTACACGACCTCTCTCTTCCACTGG |
| 9_R | TACGGTAGCAGAGACTTGGTCTAATGCCTTGTTGTCCACCTG |
| 10_F | ACACTGACGACATGGTTCTACAGGCTGCTCTCAATCACCTCT |
| 10_R | TACGGTAGCAGAGACTTGGTCTGCAGTGCTGTTAGGTGTCTC |
| 11_F | ACACTGACGACATGGTTCTACAGAGACACCTAACAGCACTGC |
| 11_R | TACGGTAGCAGAGACTTGGTCTTCCCATCAACAAGGCAGAGA |
| 12_F | ACACTGACGACATGGTTCTACAGCCATTTCAACAGCAACACC |
| 12_R | TACGGTAGCAGAGACTTGGTCTGGGGCTCTGAGATTCTTCCA |
| 13_F | ACACTGACGACATGGTTCTACAGGAAGAATCTCAGAGCCCCA |
| 13_R | TACGGTAGCAGAGACTTGGTCTCACACTAACCTGCATGCCTT |
| 14_F | ACACTGACGACATGGTTCTACAATGGACTTGTGCAAATGCCTAGTAA |
| 14_R | TACGGTAGCAGAGACTTGGTCTTGGAGGTGGACATTTATCCAACAA |
| 15_F | ACACTGACGACATGGTTCTACATGTTTTCCTGAGATCTAACCACA |
| 15_R | TACGGTAGCAGAGACTTGGTCTCAAGGCCACGTATTACTGTAACA |
| 16_F | ACACTGACGACATGGTTCTACATGTAGAACACTAAACTAGACTGCT |
| 16_R | TACGGTAGCAGAGACTTGGTCTACACAGTATTAGAAACATGCCTTTT |
| 17_F | ACACTGACGACATGGTTCTACAGTTCTGGGAGGAGGAGGAAA |
| 17_R | TACGGTAGCAGAGACTTGGTCTAGCACAGGATAACTCTTTGCA |
| 18_F | ACACTGACGACATGGTTCTACAAAACTTCCACAGGTATTTGTAGC |
| 18_R | TACGGTAGCAGAGACTTGGTCTCCAACATGGCTTAGAAGATTTCC |
| 19_F | ACACTGACGACATGGTTCTACAGTGGAAGTTGCAGCTACATGA |
| 19_R | TACGGTAGCAGAGACTTGGTCTTGCTCCCTGGAACTTTCATG |
| 20_F | ACACTGACGACATGGTTCTACAACCGTGTGCTAACCAATTGC |
| 20_R | TACGGTAGCAGAGACTTGGTCTACAAACCATTCACAGTCACCT |
| 21_F | ACACTGACGACATGGTTCTACAGGAGCTTCTTAATGTAGTTGATCC |
| 21_R | TACGGTAGCAGAGACTTGGTCTGCTGAATAAACCTATACACTGGAAC |
| 22_F | ACACTGACGACATGGTTCTACACTAATGGGTTCTTGGTGGCC |
| 22_R | TACGGTAGCAGAGACTTGGTCTTGAACCCAATGAACAGTGCC |
| 23_F | ACACTGACGACATGGTTCTACAGCTGGTCACAAATAATTTCTCCC |
| 23_R | TACGGTAGCAGAGACTTGGTCTTGAGCTGGTTCTTCTTTTGTCC |
| 24_F | ACACTGACGACATGGTTCTACAACTTGGAAAACTTTGTGGTGCT |
| 24_R | TACGGTAGCAGAGACTTGGTCTCACTGCTGCTTCATAACCCA |

### Mathematical methods

#### Calculating the crypt fusion rate

For individual crypts, fusion/fission events occur at a rate $\rho$ and have a duration $\Delta\tau$. If we take a snapshot of a piece of tissue at a time $t_0$ we see all fusion/fission events that occurred in the window $[t_0 - \Delta\tau, t_0]$. Calculating the average number of events per crypt, $X$, in a time $\Delta\tau$ over many snapshots is the same as calculating the probability of an event for a single crypt in the window $\Delta\tau$ (as we can only have a single event in any time window equal to the event duration). The number of events for a single crypt follows a Poisson distribution,

$$X_1 \sim \mathrm{Poi}(\rho\Delta\tau). \tag{0.1}$$

We want to observe events on the edge of mutant patches such that we can differentiate fission events from fusion events. For a patch with edge length $N$ (that is, $N$ crypts define the patch perimiter, each with at least one wild-type crypt as a neighbour), the number of crypts undergoing fusion or fission is distributed as

$$X_N \sim \mathrm{Poi}(N\rho\Delta\tau). \tag{0.2}$$

For a given patch with edge length $N$, then, the probability of zero events in a window $\Delta\tau$ is

$$p\{X_N = 0\} = \mathrm{e}^{-N\rho\Delta\tau} \sim 1 - N\rho\Delta\tau + \mathcal{O}(N^2\rho^2\Delta\tau^2), \tag{0.3}$$

where $N\rho$ is considered small compared with $\Delta\tau$ such that we may define a parameter $\varepsilon = N\rho\Delta\tau$ where $\varepsilon \ll 1$. Correspondingly, the probability of seeing at least one event in a window $\Delta\tau$ is

$$p\{X_N \geq 1\} = 1 - p_0 \sim N\rho\Delta\tau + \mathcal{O}(\varepsilon^2). \tag{0.4}$$

This equation can be applied to either fusion events or fission events separately by changing the event rate $\rho$ to $\rho_{\mathrm{fu}}$ or $\rho_{\mathrm{fi}}$, the event rates of fusion and fission, respectively, assuming that the event duration $\Delta\tau$ is approximately equal for fission and fusion.

#### Calculating the fusion rate

Let the number of partially mutant (partial) and fully mutant (monoclonal) fufi events observed on the edge of mutant patches be $n_p$ and $n_m$ respectively. These numbers are combined over many different mutant patches and tissue samples, with a total patch edge length of $N$. To calculate the fusion rate $\rho_{\text{fu}}$ given the fission rate $\rho_{\text{fi}}$, we use the following observations and assumptions:

1. All fission events are monoclonal.
2. Not all monoclonal events are fissions.
3. All partial events are fusions.
4. Not all fusions are partials.

The first and third assumptions stem from the belief that the timescale over which monoclonal conversion occurs in a crypt is short compared to the time between fusion/fission events. The second and fourth observations are alternative statements of the fact that fusion at the patch edge can happen inwards: a mutant crypt on the patch edge can fuse with a mutant crypt within the patch, hence creating a monoclonal event. We define the parameter $\chi$ to be the proportion of fusion events that are "inwards" and hence monoclonal. Then the ratio $n_p/n_m$ may be written as

$$\frac{n_p}{n_m} = \frac{2n_{\text{fu}}(1-\chi)}{n_{\text{fi}} + 2n_{\text{fu}}\chi}, \tag{0.5}$$

where $n_{\text{fu}}$ and $n_{\text{fi}}$ are the number of fusion and fission events, respectively. The factor of two accompanying each instance of $n_{\text{fu}}$ in (0.5) comes from the hypothesis that fusion can be initiated by either of the participating crypts, so the number of events associated with the fusion rate of any individual crypt should be half the number observed. We do not know $n_{\text{fu}}$ and $n_{\text{fi}}$ *a priori*, however.

We may recast the probability $p\{X_N \geq 1\}$ from (0.4) as the number of observed events over sample size, $n_{\text{events}}/N$, for fusion and fission, we get

$$\frac{n_{\text{fu}}}{N} \sim N\rho_{\text{fu}}\Delta\tau, \tag{0.6}$$

and

$$\frac{n_{\text{fi}}}{N} \sim N\rho_{\text{fi}}\Delta\tau, \tag{0.7}$$

where we have assumed the duration of a fusion event is approximately equal to that of a fission event. Thus, we find the approximate equivalence

$$\frac{n_{\text{fi}}}{n_{\text{fu}}} \sim \frac{\rho_{\text{fi}}}{\rho_{\text{fu}}} \tag{0.8}$$

between the ratios of numbers and rates of events. Using this, we may rewrite (0.5) as

$$\frac{n_p}{n_m} = \frac{2(1-\chi)}{\frac{\rho_{\text{fi}}}{\rho_{\text{fu}}} + 2\chi}, \tag{0.9}$$

and subsequently find an expression for the fusion rate:

$$\rho_{\text{fu}} = \frac{\rho_{\text{fi}}}{2\frac{n_m}{n_p}(1-\chi) - 2\chi}. \tag{0.10}$$

The simplest model for the parameter $\chi$ is to assume unbiased (isotropic) fusion, such that the proportion of fusions that are monoclonal is simply

$$\chi = \frac{N_t - N_w}{N_t}, \tag{0.11}$$

where $N_t$ and $N_w$ are the total number of neighbours and number of wild-type neighbours of a given fusion event, respectively.

### Diffusion model of tissue reorganisation

Below we lay out the theoretical framework and statistical methods used for understanding tissue rearrangement due to clonal expansion in the gut as a diffusion process.

#### Parameterisation and derivation

We will approach the idea of crypt packing my defining a quantity $\gamma$ that represents the local stromal fraction of the tissue (we will also refer to this as the “white space” fraction). If you look at a small region of tissue in cross-section, $\gamma$ is the fraction of that region that is taken up by stroma rather than epithelial cells (i.e. that fraction that is not part of a crypt). It is useful to think of the crypts as a density that is moving in the “free space” of the stroma. Then, we can define another quantity $\psi$ that represents this density in such a way that $\psi \equiv 1/\gamma$. This density $\psi$ tells us the number of units of area we would need to observe to see one unit of area of white space.

In intestinal tissue, we assume there is a patient-specific homeostatic degree of crypt packing that can be represented by “ambient” values $\gamma_a$, $\psi_a$ for the stromal fraction and crypt density, respectively. Near to a region of clonal expansion (a fission-driven mass source), the tissue is perturbed and the crypt density is altered such that

$$\psi(\boldsymbol{r}, t) = \psi_a + \tilde{\psi}(\boldsymbol{r}, t), \tag{0.12}$$

where the spatiotemporal variation in the crypt density is contained in the perturbation term $\tilde{\psi}(\boldsymbol{r}, t)$. By centring our polar coordinate system $\boldsymbol{r} = (r, \theta)$ at the initiation point of the clonal expansion we can state our far-field condition: $\tilde{\psi} \to 0$ as $r \to \infty$, meaning that we expect the tissue to remain in its ambient structure far from any perturbation. We choose to model the density perturbation $\tilde{\psi}$ as undergoing diffusive dynamics governed by the 2D diffusion equation

$$\frac{\partial\tilde{\psi}}{\partial t} = \nabla \cdot \left(D\nabla\tilde{\psi}\right), \tag{0.13}$$

where the coefficient of diffusion $D$ quantifies the speed with which the tissue can react to new mass being created by fission by rearranging to accommodate it. We assume $D$ is isotropic and homogeneous such that (0.13) simplifies to

$$\frac{\partial\tilde{\psi}}{\partial t} = D\nabla^2\tilde{\psi}. \tag{0.14}$$

As a way to use (0.14) to understand the dynamical process underlying observed patches of clonally-expanded mutant tissue, we note that while we know the initial size (a single crypt) and the current size ($n$ mutant crypts) of the of the clonal expansion we do not know the total age of the mutant patch nor when the individual fission events driving the expansion occurred. Thus we make the simplifying assumption that we can adequately model the system as having undergone an initial "injection" of mass equal to the change in crypt area of the clonal expansion over its lifetime. Therefore we aim to solve (0.14) with the point-source initial condition

$$\tilde{\psi}(\boldsymbol{r}, 0) = M\delta(\boldsymbol{r}), \tag{0.15}$$

where $M$ encodes the magnitude of the perturbation. We can rationalise what value $M$ should take for a given patch of $n$ crypts by the following argument: (i) we add $(n-1)$ new crypts to the initiation point of clonal expansion; (ii) as a point source perturbation to $\psi$, $M$ should equate to the extra area that we need to view in order to observe a unit of area of white space; (iii) the combined area of the $(n-1)$ new crypts will be exactly the extra area we must observe to see a unit of white space; (iv) thus $M$ should be exactly equal to the excess crypt area that the tissue must accommodate due to the clonal expansion. We may define $M$ in terms of the areas $a_{\mathrm{w}}$, $a_{\mathrm{m}}$ of wild type and mutant crypts as

$$M = (n-1)a_{\mathrm{m}} + (a_{\mathrm{m}} - a_{\mathrm{w}}), \tag{0.16}$$

where the final bracket accounts for the change in size of the crypt that gains the initial mutation.

We approach (0.14) by taking a spatial Fourier transform such that the transformed density

$$\hat{\psi}(k_x, k_y, t) = \int_{-\infty}^{\infty}\int_{-\infty}^{\infty} \tilde{\psi}(x, y, t)\mathrm{e}^{-ik_x x}\mathrm{e}^{-ik_y y}\mathrm{d}x\mathrm{d}y, \tag{0.17}$$

is governed by

$$\frac{\partial\hat{\psi}}{\partial t} = -D(k_x^2 + k_y^2)\hat{\psi}, \tag{0.18}$$

where $\boldsymbol{k} = (k_x, k_y)$ is the wave vector in a Cartesian coordinate system $(x, y)$ with its origin at the initiation point of clonal expansion. Equation (0.18) can be solved by integrating with respect to time and applying the initial condition $\hat{\psi}(0) = \mathscr{F}[M\delta(\boldsymbol{r})]$. We find

$$\hat{\psi}(\boldsymbol{k}, t) = M\mathrm{e}^{-D|\boldsymbol{k}|^2 t}. \tag{0.19}$$

We proceed by inverting the Fourier transform to find the crypt density in terms of the spatial coordinates $x$ and $y$. First, note that the inversion can be separated as follows:

$$\tilde{\psi}(x, y, t) = M\left(\frac{1}{2\pi}\right)^2 \left(\int_{-\infty}^{\infty} \mathrm{e}^{-Dk_x^2 t + ik_x x}\mathrm{d}k_x\right)\left(\int_{-\infty}^{\infty} \mathrm{e}^{-Dk_y^2 t + ik_y y}\mathrm{d}k_y\right). \tag{0.20}$$

Completing the square in the exponent of the $x$ integrand we can recast the integral as

$$\int_{-\infty}^{\infty} \mathrm{e}^{-Dk_x^2 t + ik_x x}\mathrm{d}k_x = \mathrm{e}^{-\frac{x^2}{4Dt}} \int_{u(-\infty)}^{u(\infty)} \mathrm{e}^{-Dtu^2}\mathrm{d}u, \tag{0.21}$$

where $u = k_x + ix/2Dt$ and the integral on the right hand side of (0.21) can be evaluated:

$$\int_{u(-\infty)}^{u(\infty)} \mathrm{e}^{-Dtu^2}\mathrm{d}u = \sqrt{\frac{\pi}{Dt}}. \tag{0.22}$$

By symmetry the full solution is found to be

$$\tilde{\psi}(\boldsymbol{r}, t) = \frac{M}{4\pi Dt}\mathrm{e}^{-\frac{r^2}{4Dt}}, \tag{0.23}$$

where $r^2 = x^2 + y^2$ is the radial distance from the initiation point of the clonal expansion. The stromal fraction near to a mutant patch may be expressed using (0.23) as

$$\gamma(\boldsymbol{r}, t) = \frac{1}{\psi_a + \tilde{\psi}(\boldsymbol{r}, t)} = \frac{\gamma_a}{1 + \frac{M\gamma_a}{4\pi Dt}E_d(r)}, \tag{0.24}$$

where for notational ease we have defined the frequently occurring exponential function

$$E_d(x) = \mathrm{e}^{-\frac{x^2}{4Dt}}. \tag{0.25}$$

#### Comparing theory to experiment

While (0.24) fully describes the theoretical diffusion process we posit as an explanation for the alleviation of crypt packing in the gut in lieu of mass crypt fusion, we must do more work to form a quantity suitable for comparison with experimental measurements. The data we have are measurements of the total area of patches of $n$ crypts (including their “share” of the stromal space bordering the patch) and the area of the individual crypts making up each patch. Thus we can calculate the total white space $\Gamma$ in a patch by subtracting the summed crypt areas from the total patch area. We can get a comparable theoretical quantity $\Gamma_{\mathrm{th}}$ by integrating the theoretical white space fraction (0.24) over the area $A$ occupied by a patch

$P$ of $n$ crypts:

$$\Gamma_{\mathrm{th}} = \iint_{P_A} \gamma(\boldsymbol{r}, t)\mathrm{d}\boldsymbol{A}. \tag{0.26}$$

The integral (0.26) is non-trivial for an arbitrary patch geometry. To simplify the problem we transform the geometry of the patch into a subsector of a circle centred at the coordinate origin while conserving the patch area.

To do this, three quantities must be found: the angle $\theta_s$ subtended by the subsector, its inner radius $R_{\mathrm{in}}$ and its outer radius $R_{\mathrm{out}}$. To find $\theta_s$ the distance $d$ between the centroid of the patch and the centroid of the mutant source patch (the coordinate origin) is first calculated. Then,

$$\theta_s = \begin{cases} 2\,\mathrm{atan}\dfrac{R_{\mathrm{w}}}{d}, & \text{if } d > 0, \\ 2\pi, & \text{if } d = 0, \end{cases} \tag{0.27}$$

where $R_{\mathrm{w}}$ is the radius of the patch being transformed (and where "radius" means the radius that would produce the area of the patch if the patch were a circle). Now, the area of a subsector is given by

$$A_{\mathrm{sec}} = \frac{\theta_s}{2}\left(R_{\mathrm{out}}^2 - R_{\mathrm{in}}^2\right), \tag{0.28}$$

and area conservation enforces the equality $A_{\mathrm{sec}} = \pi R_{\mathrm{w}}^2$. If $d > 0$ we may use the approximation $R_{\mathrm{in}} = d - cR_{\mathrm{w}}$ and $R_{\mathrm{out}} = d + cR_{\mathrm{w}}$ for some small $c > 0$ and substitute into (0.28) to find

$$c = \frac{\pi R_{\mathrm{w}}}{2d\theta_s}. \tag{0.29}$$

Using (0.29) to fix the value for $R_{\mathrm{in}}$ as

$$R_{\mathrm{in}} = \begin{cases} d - \dfrac{\pi R_{\mathrm{w}}^2}{2d\theta_s}, & \text{if } d > 0, \\ 0, & \text{if } d = 0, \end{cases} \tag{0.30}$$

we may then invoke area conservation once more to find

$$R_{\mathrm{out}} = \sqrt{R_{\mathrm{in}}^2 + \frac{2\pi R_{\mathrm{w}}^2}{\theta_s}}. \tag{0.31}$$

We can now approximate the integral (0.26) with

$$\Gamma_{\mathrm{th}} \approx \int_0^{\theta_s} \int_{R_{\mathrm{in}}}^{R_{\mathrm{out}}} \gamma(\boldsymbol{r}, t) r \mathrm{d}r \mathrm{d}\theta, \tag{0.32}$$

which can be evaluated to give

$$\Gamma_{\mathrm{th}} = \frac{1}{2}\theta_s \gamma_a \left\{ \left(R_{\mathrm{out}}^2 - R_{\mathrm{in}}^2\right) - 4Dt \ln\left[\frac{1 + E_d\left(R_{\mathrm{in}}\right)}{1 + E_d\left(R_{\mathrm{out}}\right)}\right] \right\}, \tag{0.33}$$

where $E_d(\cdot)$ is defined in (0.25). The theoretical value for the total patch white space defined in (0.33) can then be used to fit the diffusion model given a data set of patch measurements.

#### Statistical model for inferring the diffusion coefficient

To infer the parameters defining the tissue-intrinsic diffusion process we define the likelihood as

$$\Gamma^{(pq)} \sim \mathcal{N}\left(\Gamma_{\text{th}}^{(pq)}(D, \gamma_a^{(p)}, t^{(p)}, M^{(p)}, L^{(pq)}), \sigma_\Gamma\right), \tag{0.34}$$

where $\Gamma^{(pq)}$ and $\Gamma_{\text{th}}^{(pq)}$ are respectively the observed and theoretical total white space in patch $q$ of the neighbourhood of the $p$th mutant patch (as defined in (0.33)). The parameters defining $\Gamma_{\text{th}}^{(pq)}$ are the single tissue-intrinsic diffusion coefficient $D$, the ambient stromal fraction $\gamma_a^{(p)}$, patch age $t^{(p)}$, and input area $M^{(p)}$ of the $p$th mutant patch, and the location parameters $L^{(pq)}$ which define the transformed inner and outer radii and angle subtended by the patch $q$ of the neighbourhood of the $p$th mutant patch. We simultaneously infer the coefficient of diffusion $D$, the local ambient white spaces $\gamma_a^{(p)}$ and the patch ages $t^{(p)}$.

We use a hierarchical model to constrain $\gamma_a^{(p)}$ such that each value is drawn from a population distribution

$$\gamma_a^{(p)} \sim \text{Beta}\left(\alpha_{\text{pop}}, \beta_{\text{pop}}\right), \tag{0.35}$$

with hyperpriors

$$\alpha_{\text{pop}} \sim \text{Gamma}(10, 1), \qquad \beta_{\text{pop}} \sim \text{Gamma}(10, 1), \tag{0.36}$$

chosen to be uninformative around an unbiased mean value $\gamma_a^{\text{pop}} = 1/2$. The patch ages are measured in years and bounded in $t^{(p)} \in [5, 100]$, with priors centred on half the age of the patient $\tau^{(p)}$:

$$t^{(p)} \sim \mathcal{N}\left(\tau^{(p)}/2, \tau^{(p)}/4\right). \tag{0.37}$$

This treatment of patch ages is used to fix the parameter $t^{(p)}$ to reasonable values with the correct order of magnitude. The pairs $(D, t^{(p)})$ in isolation are unidentifiable, appearing only as a product; however, since we are fitting every mutant neighbourhood with the same diffusion coefficient we are able to infer an identifiable value for $D$ given patch ages within reasonable bounds. Standard normal distributions bounded to the positive real line are used as prior distributions for the diffusion coefficient and the standard deviation $\sigma_\Gamma$ about the theoretical total white space,

$$D \;\sim\; \mathcal{N}^{(+)}(0, 1), \qquad \sigma_\Gamma \;\sim\; \mathcal{N}^{(+)}(0, 1). \tag{0.38}$$

Inference for the full model defined by eqs. (0.34)–(0.38) was performed by MCMC sampling using the Stan probabilistic programming language.

To test whether we are justified in imposing the form of a diffusion process on the data we performed the inference above for ten shuffled data sets, where the identity of the patch responsible for the clonal expansion in each mutant neighbourhood was shuffled from the mutant clone to a random adjacent patch (and the location parameters $L^{(pq)}$ recalculated given the new structure of the neighbourhood). The output of the shuffled-data inference was assessed against the results from the raw data by two methods: (i) comparing the values of the kernel of log posterior for the MCMC sample draws (since the statistical model is identical there are no confounding dropped coefficients voiding this comparison); (ii) performing leave-one-out cross-validation on the raw log likelihood at the posterior parameter values. Both methods show that there is more evidence to support the hypothesis of identifying the mutant patch as the source of clonal expansion (that is, the inference on the raw, unshuffled data produces a better-fitting model).

#### Diffusion with a stochastically-firing point source

To investigate the temporal aspect of clonal expansion the diffusion model was extended to accommodate growth over time due to stochastic crypt fission at the patch centre. One can intuitively think of this as overlaying identical diffusion processes in space with different initiation times. Mathematically this can be achieved by breaking the solution space into chunks separated by each fission event time. If the Fourier-space density $\hat{\psi}_1(t)$ is the solution of (0.18) in a time interval $[t_0, t_1)$, achieving a value $\hat{\psi}_1(t_1)$ at the end of this period, then the solution $\hat{\psi}_2(t)$ for the next period $[t_1, t_2)$, where a fission event occurs at $t_1$, is found by solving (0.18) with the initial condition $\hat{\psi}_2(t_1) = \hat{\psi}_1(t_1) + \mathscr{F}[a_\mathrm{m}\delta(\boldsymbol{r})]$ where $a_m$ is the area of the new mutant crypt. Performing this process iteratively produces, with initial area perturbation $A_0 = a_\mathrm{m} - a_\mathrm{w}$ generated by the mutational hit, the compound diffusion solution

$$\hat{\psi}(\boldsymbol{k}, t) = A_0 \mathrm{e}^{-D|\boldsymbol{k}|^2 t} + \sum_{i=1}^{n_f} a_\mathrm{m} \mathrm{e}^{-D|\boldsymbol{k}|^2 \tau_i}, \tag{0.39}$$

where $\tau_i = \left(t - \sum_j^i t_j\right)$ for $i \in [1, n_f]$ with $t_k$ the event times of $n_f$ crypt fissions. Inverting the Fourier transform gives the full spatial solution for the density perturbation:

$$\tilde{\psi}(r, t) = \frac{A_0}{4\pi D t} \mathrm{e}^{-\frac{r^2}{4Dt}} + \sum_{i=1}^{n_f} \frac{a_m}{4\pi D \tau_i} \mathrm{e}^{-\frac{r^2}{4D\tau_i}}. \tag{0.40}$$

The formulation (0.40) supposes that we do not know how large a mutant patch will grow in a given amount of time $t$, but that we can generate fission times $t_k$ and take those events with $t_k < t$ to define our solution up to the desired time $t$. We can so generate fission times given a fission rate $\rho$ by assuming exponential waiting times (between events generated by a Poisson process) and using

$$t_n = -\frac{\ln(1 - u_{01})}{\rho n}, \tag{0.41}$$

where $n$ is the current patch size, $t_n$ is the time of the $n\mathrm{th}$ fission event and $u_{01}$ is a random draw from the unit uniform distribution $\mathcal{U}(0,1)$. So given a mutation that confers an expansion bias in terms of an increased fission rate over wild type epithelium, we can simulate the distribution of theoretical clonal expansion dynamics by generating a sequence of fission event times using (0.41) and calculate the evolution of the packing density over time using (0.40) using the diffusion coefficient inferred using the patches of ten crypts.

#### Polyp initiation

Here we quantify when the biological might break down due to physical constraints on crypt density. In the theoretical scheme defined above, as crypt density $\psi \to \infty$ the stromal fraction $\gamma \to 0$. Below some local value of $\gamma$, there will not be enough space for a crypt to fission. We posit that this may be a mechanism for outward growth, or polyp formation. A useful threshold to set on the available white space can be borrowed from the mathematics of optimal packing; identical circles may be optimally hexagonally packed to fill a fraction $\pi/\sqrt(12)$ of the space. Thus, we set the lower bound on the stromal fraction $\gamma_b = 1 - \pi/\sqrt{12}$. A mutant patch that creates mass through fission at such a rate that the tissue-intrinsic diffusion cannot act to fully accommodate new crypts will have a decrease over time of the available stromal fraction as the patch grows exponentially. We claim that if the available white space dips below $\gamma_b$ then there the patch has a finite chance of initiating outgrowth.

We can quantify this more concretely by simulating an ensemble of mutant patches with a given fission rate. At each time point we can find the radial extent $r_i$ of each instance $i$ of the mutant patch by constraining the integral over the non-stromal fraction to equal the total area of the $n_i$ mutant crypts in the patch,

$$\int_0^{2\pi}\int_0^{r_i} (1-\gamma(r))\, r\mathrm{d}r\mathrm{d}\theta = n_i a_m, \tag{0.42}$$

where the mutant crypts have area $a_m$. To approximate the effective local stromal fraction throughout the patch, the cumulative average of $\gamma(r)$, $\bar{\gamma}(r)$ is calculated starting from the patch centre $r = 0$. For each patch area $\pi r_i^2$ we may then calculate the fraction $p_f$ of this area for which $\bar{\gamma} < \gamma_b$. This gives us an idea of the portion of the patch that is at risk of initiating a polyp with the next fission event. Averaging $p_f$ over the whole ensemble, we find an estimate for the probability of a clonal expansion developing into a polyp for a given time $t$ after the initial mutation.

#### Dynamic fission-fusion as a birth-death process

The growth of clonal patches via the increase in number of clonal glands can be thought of as a birth-death process whereby crypt fission and fusion govern the dynamics of the clonal population. Fission is a perfect analogue for a birth process whereby a single gland splits to create two glands. Fusion is not a direct analogue of a death process, however; there are two possible types of fusion event: (i) fusion between two clonal glands within the patch; (ii) fusion between a gland at the patch edge and a gland in the surrounding tissue. Type (i) events will always act to reduce the clonal population by one. Type (ii) events resolve stochastically into either a wild type crypt or a patch crypt, as governed by the stem cell dynamics in the newly fused crypt base. Thus for a patch of size $n$ the "birth" rate is given by $\mu_b = n\rho_{\text{fi}}$, for the fission rate $\rho_{\text{fi}}$, and the "death" rate is given by

$$\mu_d = \rho_{\text{fi}} \left\{ (n - n_{pe}) + n_{pe}(f_{\text{in}} + f_{\text{out}}(1 - P_r)) \right\}, \tag{0.43}$$

where $n_{pe}$ is the number of crypts on the edge of the patch, $P_r$ is the bias in the stem cell competition conferred by the mutation ($P_r = 1/2$ being neutral and $P_r = 1$ being certain mutational conversion) and $f_{\text{in}}$ and $f_{\text{out}}$ are the fractions of fusion events that happen "inwards" with a within-patch crypt and "outwards" with a wild type crypts, respectively. We use the estimates

$$f_{\text{out}} = \begin{cases} 1 & \text{if } n = 1, \\ 5/6 & \text{if } n = 2, \\ 4/6 & \text{if } n \in [3, 6], \\ 1/2 & \text{otherwise,} \end{cases} \tag{0.44}$$

and $f_{\text{in}} = 1 - f_{\text{out}}$.

The value of $n_{pe}$ may be estimated from $n$ by assuming a perfect hexagonal patch shape and noting that the size of perfect hexagons may be expressed using the sum

$$n = 1 + 6 \sum_{k=1}^{L} k, \tag{0.45}$$

were $L$ is the number of "layers" around the central crypt; the patch edge length is then given by the last term of this sum ($n_{pe} = 6L$). The partial sum

$$\sum_{k=1}^{L} k = \frac{1}{2} L(L + 1) \tag{0.46}$$

is used to expand (0.45) and then the resulting quadratic in $L$ is solved to give

$$L = -\frac{1}{2} + \sqrt{\frac{1}{3}(n - 1) + \frac{1}{4}}. \tag{0.47}$$

Thus the patch edge length can be calculated as

$$n_{pe} = \begin{cases} n & \text{if } n \leq 6, \\ \text{ceil}\{6L\} & \text{otherwise.} \end{cases} \tag{0.48}$$

Instances of the resulting fission-fusion driven birth-death process were simulated using a Gillespie algorithm.
